# Conformational dynamics during misincorporation and mismatch extension defined using a DNA polymerase with a fluorescent artificial amino acid

**DOI:** 10.1101/2021.07.13.452224

**Authors:** Tyler L. Dangerfield, Serdal Kirmizialtin, Kenneth A. Johnson

## Abstract

High-fidelity DNA polymerases select the correct nucleotide over the structurally similar incorrect nucleotides with extremely high specificity while maintaining fast rates of incorporation. Previous analysis revealed the conformational dynamics and complete kinetic pathway governing correct nucleotide incorporation using a high-fidelity DNA polymerase variant containing a fluorescent unnatural amino acid. Here we extend this analysis to investigate the kinetics of nucleotide misincorporation and mismatch extension. We report the specificity constants for all possible misincorporations and characterize the conformational dynamics of the enzyme during misincorporation and mismatch extension. We present free energy profiles based on the kinetic measurements and discuss the effect of different steps on specificity. During mismatch incorporation and subsequent extension (with the correct nucleotide), the rates of the conformational change and chemistry are both greatly reduced. The nucleotide dissociation rate, however, increases to greatly exceed the rate of chemistry. To investigate the structural basis for discrimination against mismatched nucleotides, we performed all atom molecular dynamics simulations on complexes with either the correct or mismatched nucleotide bound at the polymerase active site. We show that the closed form of the enzyme with a mismatch bound is greatly destabilized due to weaker interactions with active site residues, non-ideal base pairing, and a large increase in the distance from the 3’-OH group of the primer strand to the α-phosphate of the incoming nucleotide, explaining the reduced rates of misincorporation. The observed kinetic and structural mechanisms governing nucleotide misincorporation reveal the general principles likely applicable to other high fidelity DNA polymerases.

## 1. Introduction

The molecular basis for the extraordinary specificity of enzymes has been a longstanding question in enzymology. In 1959, Koshland proposed an induced-fit model to explain enzyme specificity (1), which is a departure from the lock and key model previously proposed by Fischer (2). Koshland proposed that an enzyme active site was “not initially a negative of the substrate but became so only after interaction with substrate. This change in conformation of the protein occurred with the result that the final enzyme-substrate complex had the catalytic groups on the enzyme in the proper alignment with each other and with the bonds to be broken in the substrate molecules.” Today, structural examples of this phenomenon are abundant but the role of induced fit in enzyme specificity has been debated since it was first proposed. Specificity is a kinetic phenomenon that is difficult to resolve from structure alone. Namely, the ability of an enzyme to discriminate between competing substrates is defined by their relative *k_cat_*/*K_m_* values. Structural studies alone fail to distinguish multiple models positing how enzyme conformational changes might or might not contribute to *k_cat_*/*K_m_* values because key questions remain unanswered. How fast is the conformational change relative to the rate of the chemical reaction? Is the conformational change rapidly reversible? How do these kinetic parameters change when the enzyme encounters a different potential substrate?

DNA polymerases provide an optimal system for understanding enzyme specificity because fidelity and speed of DNA replication are biologically important for maintaining genome integrity, the alternate substrates (mismatched nucleotides as dictated by the template) are well defined, and crystal structures show large conformational changes in the enzyme after nucleotide binding (3). Moreover, measurements of free energy changes in DNA duplex formation in solution indicate relatively small differences (0.2–4 kcal/mol) between right and wrong base pairs at 37°C (4), providing a selectivity of only 5- to 100-fold which is orders of magnitude lower than the observed fidelity. Therefore, the role of the enzyme in enforcing specificity is profound. While the extraordinarily high specificity of these enzymes is well established, the roles of different steps in the kinetic pathway remains controversial (1,5–7). Of particular interest is the nucleotide-induced conformational change observed when comparing structures of binary polymerase-DNA complexes to ternary polymerase-DNA-dNTP complexes (3,8). Questions about the role of this conformational change in DNA polymerase fidelity are abundant in the literature and many theoretical arguments have been proposed (7,9,10).

The contribution of conformational changes to *k_cat_*/*K_m_* has been controversial partly due to the difficulty of measuring the rates and equilibria governing changes in enzyme structure during a single catalytic turnover. However, there is no shortage of theoretical studies speculating how conformational changes might contribute to specificity, often with conflicting ideas. The original proposal by Koshland (1) lacked any quantitative rationale for how enzyme structural changes could contribute to specificity as opposed to simply leading to increased rates of catalysis due to alignment of catalysis residues. The central question is how changes in enzyme structure can attenuate *k_cat_*/*K_m_* values for alternative substrates, not just *k_cat_* values. Herschlag attempted to delineate how conformational changes could contribute to specificity in certain cases, such as when the conformational change was rate-limiting (5). Fersht asserted that a two-step binding pathway cannot contribute more to fidelity than a thermodynamically-equivalent one-step binding (11), but his analysis assumed a rapid-equilibrium conformational change step. Post and Ray later proposed that a conformational change could still contribute to specificity when the chemical step is rate limiting but the two conformational states catalyze the reaction through different transition states (6). Tsai and Johnson provided the first evidence for a new paradigm for understanding how a fast substrate-induced conformational change could be the major determinant of specificity even when the chemistry step was rate limiting (12). Subsequent studies on HIV reverse transcriptase by Kellinger and Johnson provided additional quantitative support for this model (13,14). Countering these results, Warshel argued forcefully that pre-chemistry barriers and checkpoints cannot contribute to fidelity and catalysis as long as they are not rate limiting (7). However, Warshel failed to distinguish rate-limiting steps (*k_cat_*) from specificity-determining steps (*k_cat_*/*K_m_*), which need not be identical.

Not all polymerases follow the Tsai-Johnson induced-fit paradigm. In particular, for low fidelity enzymes Pol β and Klenow, the pre-chemistry conformational changes appear to be rapid-equilibrium steps coupled to nucleotide binding (15–18). HIV reverse transcriptase may be an exception to the rule because it has only moderate fidelity and must replicate both RNA and DNA. In addition, there are concerns over the validity of the conclusions in the original Tsai-Johnson paper because mutations required to make a cys-light variant to site-specifically label the enzyme altered its fidelity. Therefore, it is especially important to rigorously test the role of enzyme conformational dynamics in specificity using a high-fidelity polymerase that does not require construction of a cys-light variant.

The T7 DNA polymerase has long been an important, simple model system for understanding fidelity as only two polypeptides (T7 gene product 5 and thioredoxin) are required for robust polymerization and exonuclease proofreading activity (3,9,19–29). We previously reported methods to site specifically incorporate the fluorescent unnatural amino acid (7-hydroxy-4-coumarin-yl)-ethylglycine (7-HCou) into the fingers domain of T7 DNA polymerase. We showed that this variant has similar fidelity to the wild-type enzyme, while the fluorescent amino acid gives a signal to measure the nucleotide-induced conformational change (30). We determined the complete kinetic pathway of correct nucleotide incorporation, including measurements of the rates of translocation during processive synthesis (31). Full understanding of DNA polymerase fidelity, however, requires parallel analysis of the pathway of misincorporation, including the role of conformational changes, which we address in this paper. Here we examine the kinetics of mismatch formation using an exonuclease-deficient form of the enzyme (D5A/E7A) to measure the innate fidelity during the DNA polymerization step (32). In subsequent studies, we examine the kinetic basis for selectivity of the proofreading exonuclease to assess its role in overall fidelity.

To obtain an estimate for the fidelity for the T7 DNA polymerase, we began by measuring specificity constants (*k_cat_*/*K_m_*) for each possible misincorporation and then determined the discrimination against each mismatch to calculate the overall range of fidelity values for the enzyme. We then determined the complete kinetic pathway for T:dTTP misincorporation and extension of this mismatch with dTTP (when the next templating base is an A), including measurement of the dynamics of conformational changes along the pathway. To gain structural insights for bound mismatches at the polymerase active site, we performed all atom molecular dynamics (MD) simulations on three complexes and report the resulting active site structures, along with quantification of hydrogen bond parameters as well as distances from the 3’-OH of the primer to the α-phosphate of the incoming nucleotide. These structures highlight the misalignment of catalytic residues and the non-canonical base pairing that occurs with mismatches in the active site of the polymerase.

## 2. Results

### 2.1. Mismatch discrimination range for T7 DNA polymerase

While *k_cat_*/*K_m_* values for a correct- and a mismatched-nucleotide incorporation have been reported (9,29–31,33,34), general questions about misincorporation still remain, especially for high fidelity DNA polymerases. Are certain mismatches incorporated more efficiently than others? What is the range of discrimination against different mismatch combinations? To address these questions, we performed single turnover kinetic experiments with all 12 possible mismatch combinations to estimate *k_cat_*/*K_m_* for each. Measurements of the specificity constant (*k_cat_*/*K_m_*) were used to calculate a discrimination index (D) against each mismatch defined by the ratio of *k_cat_/K_m_* values for correct (measured previously (33)) versus mismatched nucleotide incorporation.

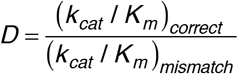

In our single turnover kinetic experiments, a 27 nt 5’-[6-FAM] (6-carboxyfluorescein) labeled primer was annealed to a 45 nt template containing either A, C, G, or T as the templating base at position 18 (see Table 1 and Figure 1). Since the experiments were performed with an excess of enzyme over DNA, the observed time dependence of product formation defined the rate of reaction at the active site of the enzyme during a single turnover. Various nucleotide concentrations were mixed with the enzyme-DNA complex to start the reaction, then samples were quenched by adding EDTA and products were resolved and quantified by capillary electrophoresis (see Materials and Methods).

**Table 1:**
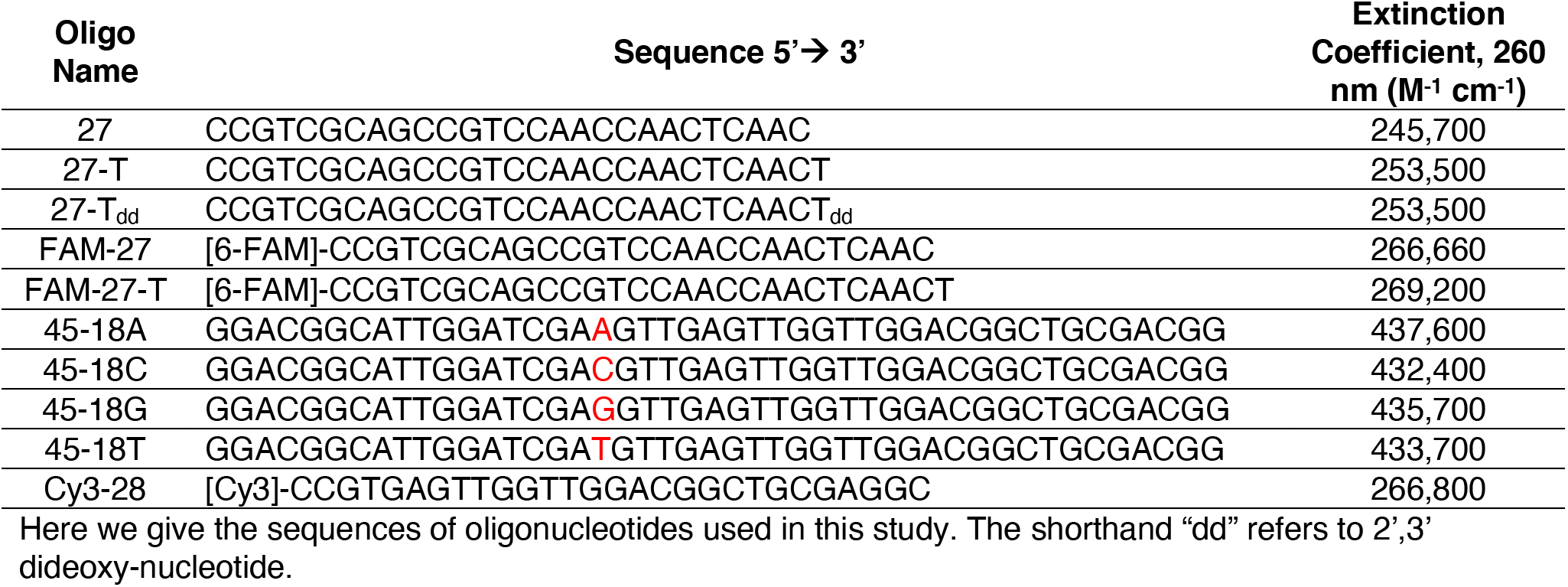
Oligonucleotides used in this study.

**Figure 1:**
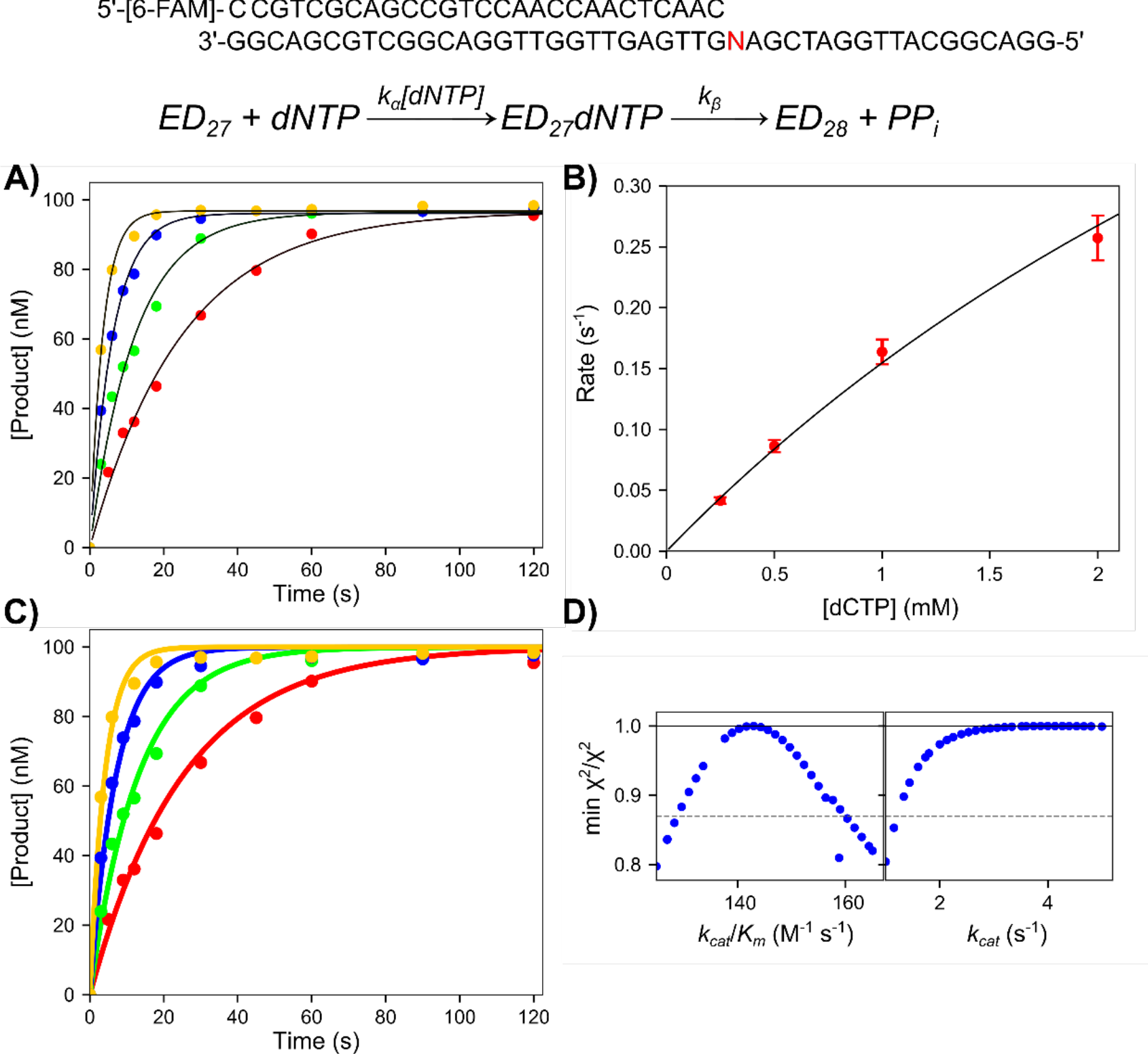
Representative mismatch incorporation kinetics. Reaction conditions: A solution of 250 nM T7 DNA polymerase E514Cou, 100 nM FAM-27/45-18N DNA (where N is the templating base for the incoming nucleotide), 5 μM thioredoxin, and 0.1 mg/ml BSA was mixed with 0.25–2 mM Mg^2+^dNTP and 12.5 mM Mg^2+^ to start the reaction. Reactions were quenched with EDTA. <u>DNA substrate:</u> The DNA substrate shown was used in the mismatch screening experiments and consists of a 27 nt 5’-6-FAM labeled primer, annealed to a 45 nt template, where the N can be A, C, G or T depending on the misincorporation reaction. <u>Scheme</u>: Simplified model for nucleotide incorporation to derive *k_cat_*/*K_m_* and *k_cat_*. The rate constant *k_α_* in this model is *k_cat_*/*K_m_* and *k_β_* is *k_cat_*. This numbering scheme was adopted to avoid confusion with more complete models shown later. <u>A) Plot of product concentration versus time for the A:dCTP misincorporation reaction</u>. The data were fit to a single exponential function (black lines). <u>B) Rate versus concentration plot for A:dCTP misincorporation reaction.</u> Observed rates are from the single exponential fits to the data in (A). The data are shown fit to a hyperbola. At the concentrations tested, there was little curvature in the rate versus concentration plot, indicating that *k_cat_*/*K_m_* (defined by the initial slope) is well defined by the data, but *k_cat_* and *K_m_* individually are not well defined. <u>C) Sample plot of product versus time for the A:dCTP misincorporation reaction–fit by simulation.</u> The data are the same as shown in (A), however the solid-colored lines through the data are the best fit by simulation in KinTek Explorer to the simple scheme at the top of the figure. <u>D) Sample confidence contours for A:dCTP misincorporation reaction.</u> Confidence contours were generated with the FitSpace function of KinTek Explorer to extract *k_cat_*/*K_m_* and a lower limit on *k_cat_*. The results from this analysis for all mismatch combinations are summarized in Table 2.

To designate various base-pair combinations, we use the shorthand notation X:dNTP where X is the templating base and dNTP is the incoming nucleotide. The A:dCTP misincorporation reaction was chosen as a representative data set to illustrate the results and methods of analysis as shown in Figure 1. Initially data were analyzed by conventional methods by fitting each reaction time course to a single exponential function (Figure 1A), then plotting the observed rate versus nucleotide concentration (Figure 1B). The rate versus concentration plot was then fit to a hyperbolic function to estimate *k_cat_*/*K_m_*. Since mismatch binding is weak, as demonstrated by the lack of significant curvature in the rate versus concentration plot, we cannot extract accurate values for *k_cat_* and *K_m_* individually, but *k_cat_/K_m_* is well defined from the initial slope of the concentration dependence of the observed rate. Fitting the data by simulation (Figure 1C) in KinTek Explorer (35,36) allows lower limits on *k_cat_* and *K_m_* to be determined in addition to providing accurate estimates for *k_cat_*/*K_m_*.(37,38) The data were fit using the kinetic model at the top of Figure 1, where *k_α_* = *k_cat_*/*K_m_* and *k_β_* = *k_cat_*, respectively. We use this nomenclature rather than numbered rate constants to avoid confusion later on when we relate these data to a comprehensive model.

We used the FitSpace function (39) of KinTek Explorer to evaluate confidence limits for each of these parameters, showing a well constrained value for *k_cat_*/*K_m_* and setting a lower limit on *k_cat_* (Figure 1D). This analysis was repeated for all possible mismatch combinations to give the results summarized in Table 2. A threshold in *χ*^2^_min_/*χ*^2^ equal to 0.85, (based on the number of parameters and data points in the fitting, calculated by the software using the F-distribution) was used to estimate the 95% confidence interval for each parameter. The data for the A:dGTP mismatch was the only case where a well-defined value for *k_cat_*/*K_m_* was not obtained from the data fit because of the biphasic nature of the data that did not fit our simple model. We therefore performed traditional equation-based fitting of the data to get an approximate estimate of *k_cat_*/*K_m_* for this mismatch (Table 2), although we caution that there are likely systematic errors on this value because the single exponential fits do not accurately describe the data. Data obtained for this mismatch are shown in Figure S1.

**Table 2:** Kinetic parameters for all possible templating base/dNTP combinations.

| Templating Base | dNTP | $k_{cat}/K_m$ (M<sup>-1</sup> s<sup>-1</sup>) | $k_{cat}$ (s<sup>-1</sup>) | $K_m$ (mM) | Discrimination |
| --- | --- | --- | --- | --- | --- |
| A | dATP | 147 ± 20 | ≥ 1.3 | ≥ 8.8 | 143,500 ± 31,200 |
|  | dCTP | 143 ± 15 | ≥ 1.2 | ≥ 8.4 | 144,200 ± 28,800 |
|  | dGTP* | 450 * | * | * | * |
|  | dTTP | [1.65 ± 0.28] × 10<sup>7</sup> | 297 ± 10 | 0.018 ± 0.003 | 1 |
| C | dATP | 190 ± 25 | ≥ 0.4 | ≥ 2.1 | 92,000 ± 20,000 |
|  | dCTP | 1.4 ± 0.2 | ≥ 0.075 | ≥ 50 | 12,600,000 ± 2,800,000 |
|  | dGTP | [1.37 ± 0.24] × 10<sup>7</sup> | 246 ± 12 | 0.018 ± 0.003 | 1 |
|  | dTTP | 31 ± 4 | ≥ 0.6 | ≥ 19 | 563,000 ± 122,000 |
| G | dATP | 295 ± 40 | ≥ 0.7 | ≥ 2.4 | 110,000 ± 21,000 |
| | dCTP | $[2.63 \pm 0.36] \times 10^7$ | 447 ± 30 | 0.017 ± 0.002 | 1 |
|  | dGTP | 61 ± 8 | ≥ 0.3 | ≥ 4.9 | 531,000 ± 100,000 |
|  | dTTP | 195 ± 20 | ≥ 1.1 | ≥ 5.6 | 162,000 ± 28,000 |
| T | dATP | $[1.56 \pm 0.26] \times 10^7$ | 280 ± 4 | 0.018 ± 0.003 | 1 |
|  | dCTP | 43 ± 5 | ≥ 2.1 | ≥ 48 | 454,000 ± 93,000 |
|  | dGTP | 265 ± 40 | ≥ 1.3 | ≥ 4.9 | 76,000 ± 17,000 |
|  | dTTP | 130 ± 25 | ≥ 0.5 | ≥ 3.8 | 161,000 ± 41,000 |

\* The A:dGTP misincorporation data were biphasic but did not fit any simple model well enough to derive $k_{cat}$ and $K_m$ . The estimate presented here is from the conventional fitting of the data to exponential functions although there is likely a large error on this value. The data are shown in Figure S1.

From the measured *k_cat_*/*K_m_* values for each mismatched base pair from this study and previously reported *k_cat_*/*K_m_* values for all correct nucleotide incorporations (33), we calculated the discrimination against each mismatch which can also be thought of as the average number of bases incorporated before an error is made. The highest discrimination index was for the C:dCTP mismatch, with a *k_cat_*/*K_m_* of 1.4 ± 0.2 M^−1^s^−1^ and a discrimination index of 12,600,000. The lowest discrimination was for the T:dGTP mismatch, with a *k_cat_*/*K_m_* of 265 ± 40 M^−1^s^−1^ and a discrimination index of 76,000. The average discrimination index was 220,000, indicating that the T7 DNA polymerase replicated DNA with high fidelity in the absence of the proofreading exonuclease. We have not extended these measurements to look for sequence context effects, but rather consider our results to represent a reasonable estimate of fidelity with smaller effects introduced by sequence context effects such as base stacking interactions (40,41).

### 2.2. T:dTTP misincorporation and mismatch extension experiments

To fully understand the role of induced-fit in DNA polymerase fidelity, we need to compare the kinetics of conformational changes during misincorporation with those during correct incorporation (31). In our previous work, we found that for correct nucleotide incorporation, all nucleotides gave a fluorescence signal that reflected changes in enzyme structure before and after incorporation (30). In this study, we screened all misincorporation combinations for a signal in the stopped-flow instrument (data not shown) and found that most combinations failed to give a usable signal. This observation is consistent with our prior interpretation of mismatch incorporation by HIVRT where the reverse of the conformational change leading to release of a mismatched nucleotide was so fast that no signal could be observed (42). In one case, with a T:dTTP misincorporation we observed a good fluorescence signal. Note that after T:dTTP misincorporation, the next templating base is an A (see DNA substrate in Figure 2) so this mismatch gets extended with the correct base although no further extension past the 29 nt product was observed. Fortuitously, the signals associated with the correct incorporation occurring after the mismatch provided unique information to define the kinetics conformational changes during misincorporation. To better estimate the kinetic parameters *k_cat_* and *K_m_* for this particular reaction and to provide a solid foundation for interpretation of stopped-flow fluorescence data, chemical-quench experiments were performed at multiple nucleotide concentrations using two protocols. In on we monitored the two sequential incorporations (misincorporation followed by mismatch extension reaction; 27 nt to 29 nt) in a single reaction. In a separate experiment we only measured the kinetics of the mismatch extension reaction by starting with a preformed mismatch (28 nt to 29 nt). The two experiments could then be fit globally.

**Figure 2:**
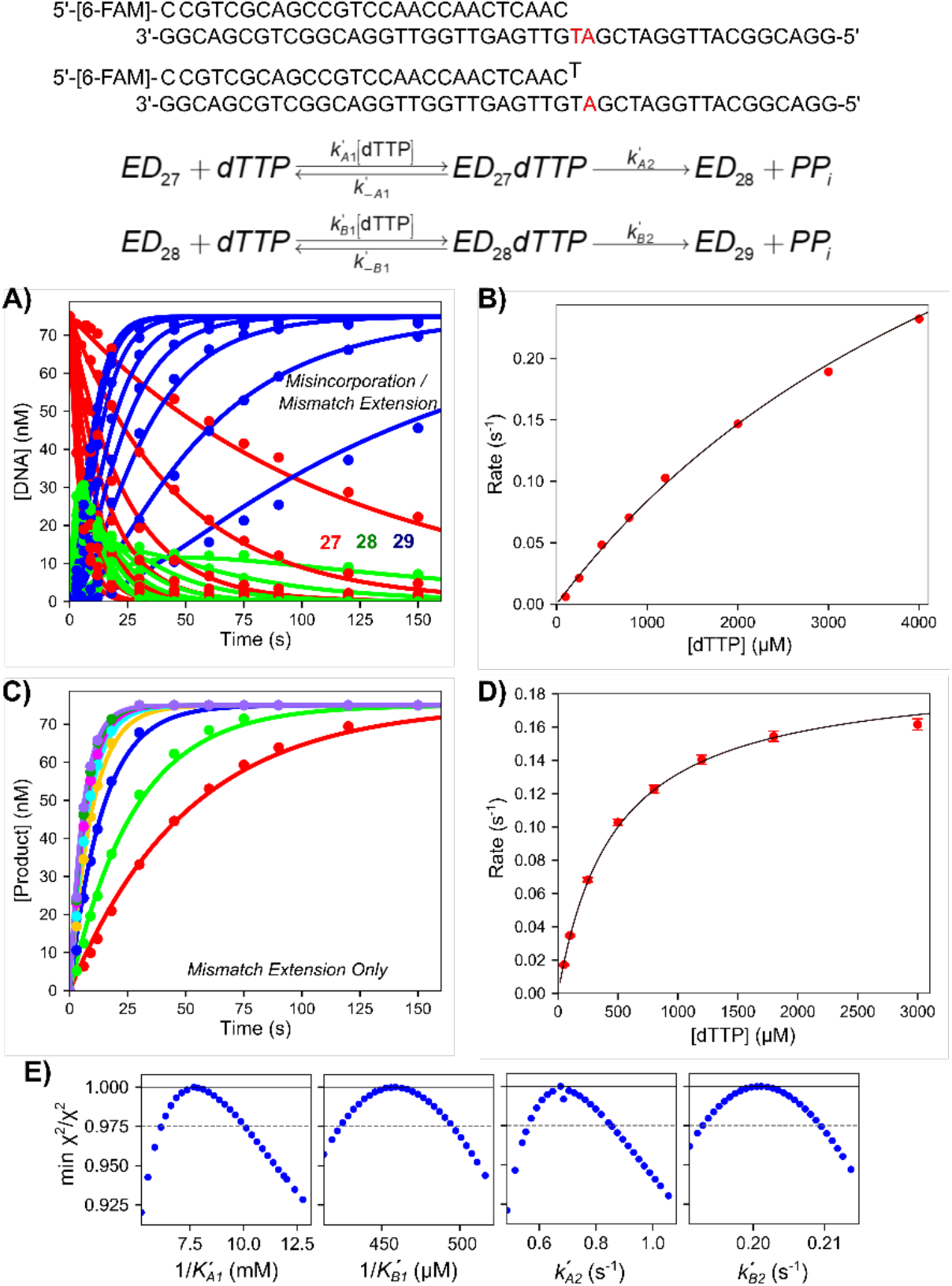
Chemical-quench dTTP misincorporation and mismatch extension experiments. Reaction conditions: A solution of 225 nM T7 DNA polymerase E514Cou, 75 nM FAM-DNA (FAM-27/45-18T or FAM-27-T/45-18T), 4.5 μM thioredoxin, and 0.1 mg/ml BSA was mixed with Mg^2+^-dTTP (0.1–4 mM for misincorporation, 0.05–3 mM for mismatch extension) and 12.5 mM Mg^2+^ to start the reaction. Reactions were quenched with EDTA. <u>DNA Substrates:</u> The top DNA substrate (FAM-27/45-18T) was used in the experiment in (A). The second DNA substrate (FAM-27-T/45-18T) was used in the experiment in (C). <u>Scheme:</u> Kinetic scheme for misincorporation followed by mismatch extension. Rate constants for the misincorporation reaction are denoted with a subscript *An*, where *n* is the step number. Rate constants for the mismatch extension are given with a subscript *Bn*, where *n* is the step number. <u>A) Time course of misincorporation and mismatch extension.</u> Various concentrations of dTTP were mixed with the enzyme-DNA 27/45 complex and the time course of misincorporation to form the 28 nt product, and subsequent mismatch extension to the 29 nt product are shown. The solid lines through the data are the best fits by simulation to the scheme at the top of the figure. The 27 nt starting material is shown in red, the 28 nt misincorporation product is shown in green, and the final 29 nt product is shown in blue. <u>B) Rate versus dTTP concentration for the misincorporation reaction.</u> Rates are from single exponential fits to the data for the loss of the 27 nt starting material (not shown) in (A). Data are shown fit to a hyperbola, and the best fit parameters from the fit are: maximum rate: 0.58 ± 0.10 s^−1^, *K_d,app_* = 6 ± 1.3 mM. <u>C) Time course of mismatch extension.</u> Various concentrations of dTTP were mixed with the enzyme-DNA 27-T/45 complex and the time course of extension to the 29 nt product is shown. Solid lines through the data are best fits by simulation in KinTek Explorer. <u>D) Rate versus dTTP</u> <u>concentration for the mismatch extension reaction.</u> Rates are from single exponential fits to the data for the formation of the 29 nt product (not shown) from the data in (C). Data are shown fit to a hyperbola, and the best fit parameters from the fit are: maximum rate: 0.194 ± 0.005 s^−1^, *K_d,app_* = 470 ± 22 μM. <u>E) Confidence contours for misincorporation and mismatch extension</u> <u>reaction.</u> Confidence contours were derived from global fitting of both data sets shown in this figure in (A) and (C) by simulation in KinTek Explorer. The grey dashed line represents the *χ*^2^ threshold to give the 95% confidence interval for each parameter (reported in Table 3).

Figure 2A shows the full reaction sequence involving misincorporation followed by mismatch extension. A solution of T7 DNA polymerase E514Cou-DNA complex (upper DNA substrate in Figure 2) was mixed with 0.1–4 mM Mg^2+^-dTTP and 12.5 mM excess Mg^2+^, then samples were quenched at various times by adding EDTA. Conventional data analysis was performed by fitting the loss of the 27 nt starting material using a single exponential function (not shown), then plotting the observed rate versus dTTP concentration (Figure 2B). The rate versus concentration plot was then fit to a hyperbola to estimate the maximum rate of 0.58 ± 0.10 s^−1^, and a *K_m_*: 6.0 ± 1.3 mM for the misincorporation reaction. Figure 2C shows the results of a separate experiment where the mismatch extension reaction was monitored by mixing the preformed enzyme-DNA complex (lower DNA in Figure 2, containing a terminal T:T mismatch) with Mg^2+^-dTTP and Mg^2+^. No further extension past the 29 nt product was observed in either experiment on the timescales examined. For conventional fitting, these data were fit to a single exponential function, then the observed rate was plotted as a function of dTTP concentration (Figure 2D). The hyperbolic fit to the rate versus concentration data estimated a maximum rate of 0.194 ± 0.005 s^−1^ and a *K_m_* of 470 ± 22 μM for the mismatch extension reaction.

Global data fitting of the two experiments simultaneously was then performed using the kinetic model shown at the top of Figure 2. Both data sets could be fit globally using the single model, indicating that there are no kinetically significant steps occurring between misincorporation and mismatch extension. The extent to which parameters of the model are defined by the data was examined using confidence contour analysis to give the results shown in Figure 2E. All confidence contours were approximately parabolic, indicating that each parameter was well constrained by the data with well-defined lower and upper limits. The best fit value and 95% confidence interval for each parameter are given in Table 3 and agree with the results from conventional equation-based data fitting. Comparing Figures 2B and 2D shows that the apparent binding affinity of dTTP for the mismatch extension reaction is much greater than for binding of dTTP for the misincorporation reaction. However, the maximum rate for the misincorporation reaction is much faster than the maximum rate for the mismatch extension reaction. This phenomenon was previously shown for this enzyme for extension on top of an A:A mismatch with dCTP as the next correct base (9). To fully understand the kinetic basis for this result, further experiments were performed to probe the role of conformational dynamics in the mismatch extension reaction.

**Table 3:**
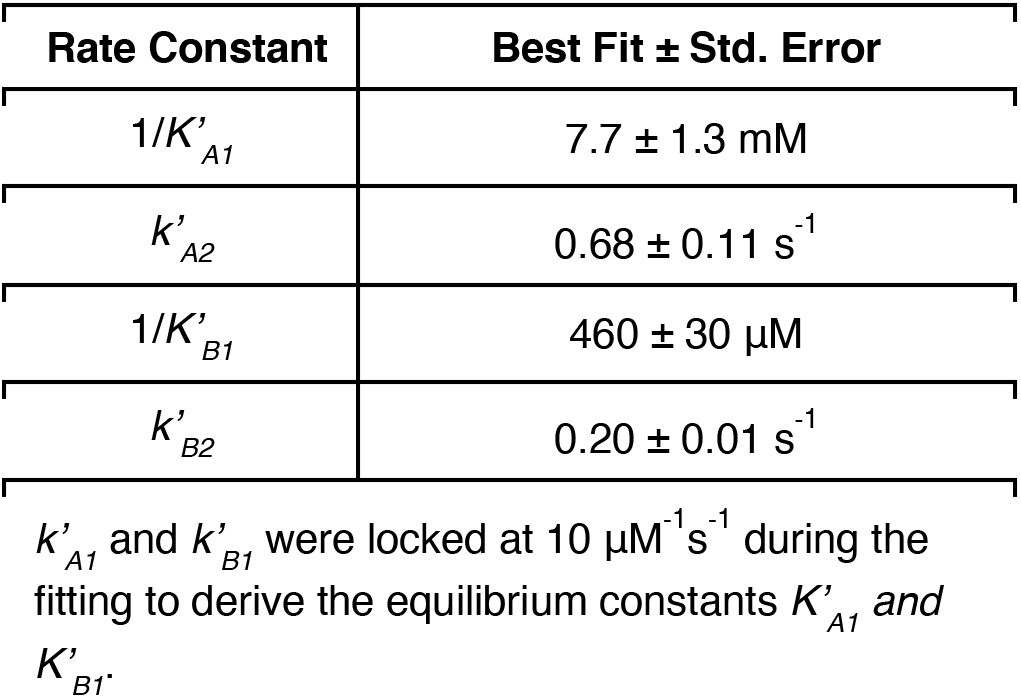
Rate constants from misincorporation and mismatch extension experiments.

| Rate Constant | Best Fit ± Std. Error |
| --- | --- |
| $1/K'_{A1}$ | $7.7 \pm 1.3$ mM |
| $k'_{A2}$ | $0.68 \pm 0.11$ s<sup>-1</sup> |
| $1/K'_{B1}$ | $460 \pm 30$ μM |
| $k'_{B2}$ | $0.20 \pm 0.01$ s<sup>-1</sup> |

$k'_{A1}$ and $k'_{B1}$ were locked at $10 \mu\text{M}^{-1}\text{s}^{-1}$ during the fitting to derive the equilibrium constants $K'_{A1}$ and $K'_{B1}$ .

### 2.3. Kinetics of nucleotide binding for correct dTTP extension on a T:T mismatch

Monitoring the complete dTTP misincorporation/mismatch extension reaction in the stopped-flow is complex, as we will later show. Therefore, we begin by measuring the conformational changes during nucleotide binding for the mismatch extension reaction only, using a DNA substrate with a 2’,3’dideoxy terminated primer to limit the observed kinetics to nucleotide binding steps (DNA substrate in Figure 3). We first measured the kinetics of nucleotide binding in the stopped-flow instrument (Figure 3A). A solution of the T7 DNA polymerase E514Cou-DNAdd complex was mixed with 0.04–3 mM Mg^2+^-dTTP and 12.5 mM Mg^2+^. The data at all concentrations fit well to a single exponential function (not shown) and revealed an increase in fluorescence upon nucleotide binding, which is a measure of nucleotide-induced enzyme closure as shown for correct nucleotide incorporation (30). To aid in visualization of the underlying mechanism, the rate versus dTTP concentration plot is shown in Figure 3B and fit to a hyperbola. The y-intercept of this plot estimates the nucleotide dissociation rate (*k_-B2_* in the scheme in Figure 3) of 5.8 ± 0.70 s^−1^, a maximum rate (approximating *k_B2_*) of 330 ± 16 s^−1^, and an apparent *K_d_* (approximating 1/*K_B1_*) of 1.56 ± 0.154 mM.

**Figure 3:**
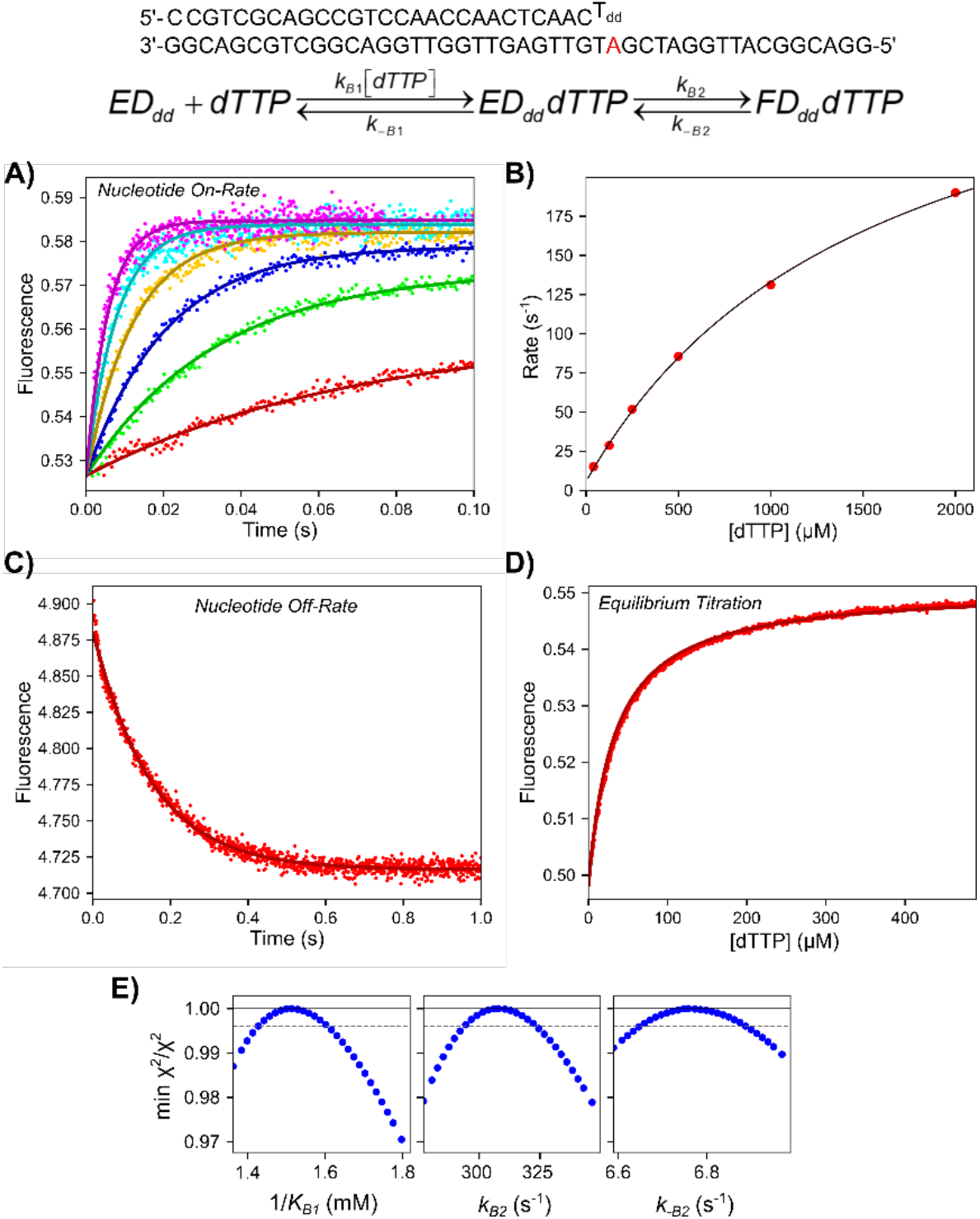
dTTP binding kinetics for mismatch extension reaction. <u>DNA Substrate:</u> The DNA substrate used in the experiments in this figure is shown. The primer strand contains a 2’,3’ dideoxyT (Tdd) as the terminal base to limit the observed kinetics to nucleotide binding steps. <u>Scheme:</u> The kinetic model for two-step dTTP binding used in data fitting is shown, where initial ground state binding (*K_B1_*, *k_−B1_*) is followed by an isomerization to a closed state (*k_B2_*, *k_−B2_*). <u>A) Stopped-flow dTTP binding-</u> <u>rate experiment.</u> A solution of 200 nM T7 DNA polymerase E514Cou, 4 μM thioredoxin, and 250 nM DNAdd (27-Tdd/45-18T) was mixed with 0.05–3 mM Mg^2+^-dTTP and 12.5 mM Mg^2+^ to start the reaction. Solid lines through the data are the best global fit by simulation to the dTTP binding data in this figure. <u>B) Rate versus concentration for dTTP on-rate experiment.</u> Rates are from single exponential fits to the data in (A). The data are shown fit to a hyperbola to extract the following parameters: y-intercept: 5.8 ± 1.7 s^−1^, *k_max_* = 332 ± 18 s^−1^, *K_d,app_* = 1560 ± 154 μM. <u>C) Stopped-flow dTTP dissociation-rate experiment.</u> A solution of 1 μM T7 DNA polymerase E514Cou, 1 μM DNAdd (27-Tdd/45-18T), 5 μM dTTP, 20 μM thioredoxin, and 12.5 mM Mg2+ was mixed with 20 μM wild-type (exo-) T7 DNA polymerase, 20 μM DNA 27/45-18A, and 20 μM thioredoxin as a trap for nucleotide released from the labeled enzyme-DNAdd-dTTP complex. The solid line shows the best global fit by simulation including the other experiments in this figure. Fitting the trace to a single exponential function (not shown) gives an observed rate of 6.6 ± 0.1 s^−1^. <u>D) Equilibrium dTTP binding fluorescence titration.</u> A solution of 100 nM T7 DNA polymerase E514Cou, 2 μM thioredoxin, 150 nM DNAdd (27-Tdd/45-18T), and 12.5 mM Mg2+ was preincubated in a cuvette in a temperature-controlled holder on the TMX module for the KinTek SF-300X instrument. A solution of 7.5 mM Mg^2+^-dTTP was titrated into the cuvette from a Hamilton syringe over the course of 5 minutes with constant stirring. The fluorescence signal was corrected for the inner filter effect and for the small dilution during the titration. The solid line through the data is the best global fit by simulation including the other data in this figure. Fitting the titration to a hyperbola (not shown) gives a *K_d_* = 37 ± 1 μM. <u>E) Confidence contours from dTTP mismatch extension binding data.</u> The dashed line gives the *χ*^2^ threshold defining the 95% confidence interval. Values from the confidence contour analysis can be found in Table 4.

To directly measure the nucleotide dissociation rate, a preformed T7 DNA polymerase E514Cou-DNA_dd_-dTTP complex was mixed with a large excess of wild-type T7 DNA polymerase-DNA complex (27/45-18A) as a trap for nucleotide released from the ternary labeled enzyme complex (Figure 3C). The data fit a single exponential function (not shown), with an observed rate of 6.6 ± 0.1 s^−1^, consistent with y-intercept from the on-rate measurement in Figure 3B, defining the rate constant for nucleotide release. Finally, an equilibrium titration of dTTP into the T7 DNA polymerase E514Cou-DNA_dd_ complex was performed. A solution of T7 DNA polymerase E514Cou-DNA_dd_ complex in a cuvette was incubated in a temperature-controlled cuvette holder on the titration module for the stopped-flow instrument. A solution of Mg^2+^-dTTP was titrated into the cuvette from a Hamilton syringe over the course of 5 minutes with constant stirring from a micro stir bar. The fluorescence intensities were corrected for the small dilution during the titration, then further corrected for the inner filter effect as described in the Materials and Methods section. The data fit a hyperbola (not shown), estimating a *K_d_* of 37 ± 1 μM. All the data in Figure 3 were then globally fit in KinTek Explorer using the model in the figure. The best fit is represented by the solid lines superimposed on the data. Confidence contour analysis shows that all parameters are well constrained by the data (Figure 3E) with best fit values summarized in Table 4.

**Table 4:** Rate constants from pre-chemistry mismatch extension experiments.

| Rate Constant | Best Fit ± Std. Error |
| --- | --- |
| $1/K_{B1}$ | $1.5 \pm 0.09$ mM |
| $k_{B2}$ | $310 \pm 14$ s<sup>-1</sup> |
| $k_{-B2}$ | $6.8 \pm 0.13$ s<sup>-1</sup> |

\* $k_{B1}$ was not individually defined by the data so this rate constant was locked at $10 \mu\text{M}^{-1}\text{s}^{-1}$ in the fitting to derive the equilibrium constant $K_{B1}$ .

While the data in Figure 3 give a simple model for nucleotide binding preceding the mismatch extension reaction, the rate constants from the stopped-flow experiments are not consistent with the chemical-quench data presented in Figure 2C. Namely, the *K_m_* from the chemical-quench experiments is at least 10-fold higher than the binding affinity measured in the stopped-flow experiments. With a maximum rate of approximately 0.2 s^−1^ and a nucleotide dissociation rate of around 7 s^−1^, the binding should reach equilibrium and the *K_d_* from the equilibrium titration in Figure 3D should match the *K_m_* from the experiment in Figure 2C. This discrepancy could be due to an artifact introduced by used of the 2’,3’-dideoxy terminated primer of the DNA substrate or there could be some other mechanism to reconcile the two data sets. As an alternative approach to using dideoxy-terminated DNA, we attempted to use the non-hydrolysable nucleoside analog dTpNpp with DNA containing a normal 3’-OH group to measure nucleotide binding kinetics; however, we found that this analog was a very poor substrate for T7 DNA polymerase (data not shown). Instead, we next performed a stopped-flow experiment with normal dTTP and DNA containing a 3’-OH to compare the stopped-flow fluorescence kinetics with the chemical-quench data and the binding data given in Figure 3.

### 2.4. Kinetics of nucleotide incorporation for correct dTTP extension on a T:T mismatch

A stopped-flow experiment was performed to directly measure the conformational changes during mismatch extension to compare with the chemical-quench experiment. A solution of preformed T7 DNA polymerase E514Cou-DNA (27-T/45-18T, primer/template shown in Figure 4) complex was mixed with 0.05–3 mM Mg^2+^-dTTP and 12.5 mM Mg^2+^ to start the reaction in the stopped-flow (Figures 4A and 4B). As with the experiments with the 2’,3’ dideoxy DNA substrate, the data show an increase in fluorescence upon nucleotide binding (shown more clearly on a log timescale in Figure 4B), but also show a slow decrease to return to the starting fluorescence level on a longer timescale. The data fit 3 exponentials at concentrations up to 750 μM and at least 4 exponentials at higher nucleotide concentrations (not shown). For conventional fitting of the data to aid in visualizing the underlying mechanism, the traces were fit to a 3-exponential function (not shown). Rates for the fast phase versus dTTP concentration are given in Figure 4C, fit to a hyperbola to extract the y-intercept: 5.9 ± 0.32 s^−1^, amplitude: 54 ± 2 s^−^1, and apparent *K_d_* = 1.2 ± 0.12 mM. Compared to the data with the dideoxy primer, the y-intercept and apparent *K_d_* are consistent, although the maximum rate was somewhat lower for the 3’-OH DNA substrate. The second and third phases are shown in Figure 4D at dTTP concentrations only up to 750 μM, since at higher concentrations there are additional slower phases. The maximum rates for the 2^nd^ and 3^rd^ phase from this fitting were 0.61 ± 0.06 s^−1^ and 0.062 ± 0.001 s^−1^, respectively. Since the maximum rate obtained from the chemical-quench experiment in Figure 2C was around 0.2 s^−1^, the chemistry step must occur between the 2^nd^ and 3^rd^ phases of the stopped-flow data. These data suggest a 3-step binding mechanism where each state has a different fluorescence scaling factor. There are numerous problems involved with conventional fitting of fluorescence data using a 3-exponential function because the function includes seven unknowns and inherent relationships between rates and amplitudes are lost. These issues can be resolved using fitting by simulation, including rapid quench data to aid in the interpretation of the fluorescence signals. Fitting the data globally uses far fewer unknown parameters and can establish the relationships between fluorescence changes and chemical reaction steps (38).

**Figure 4:**
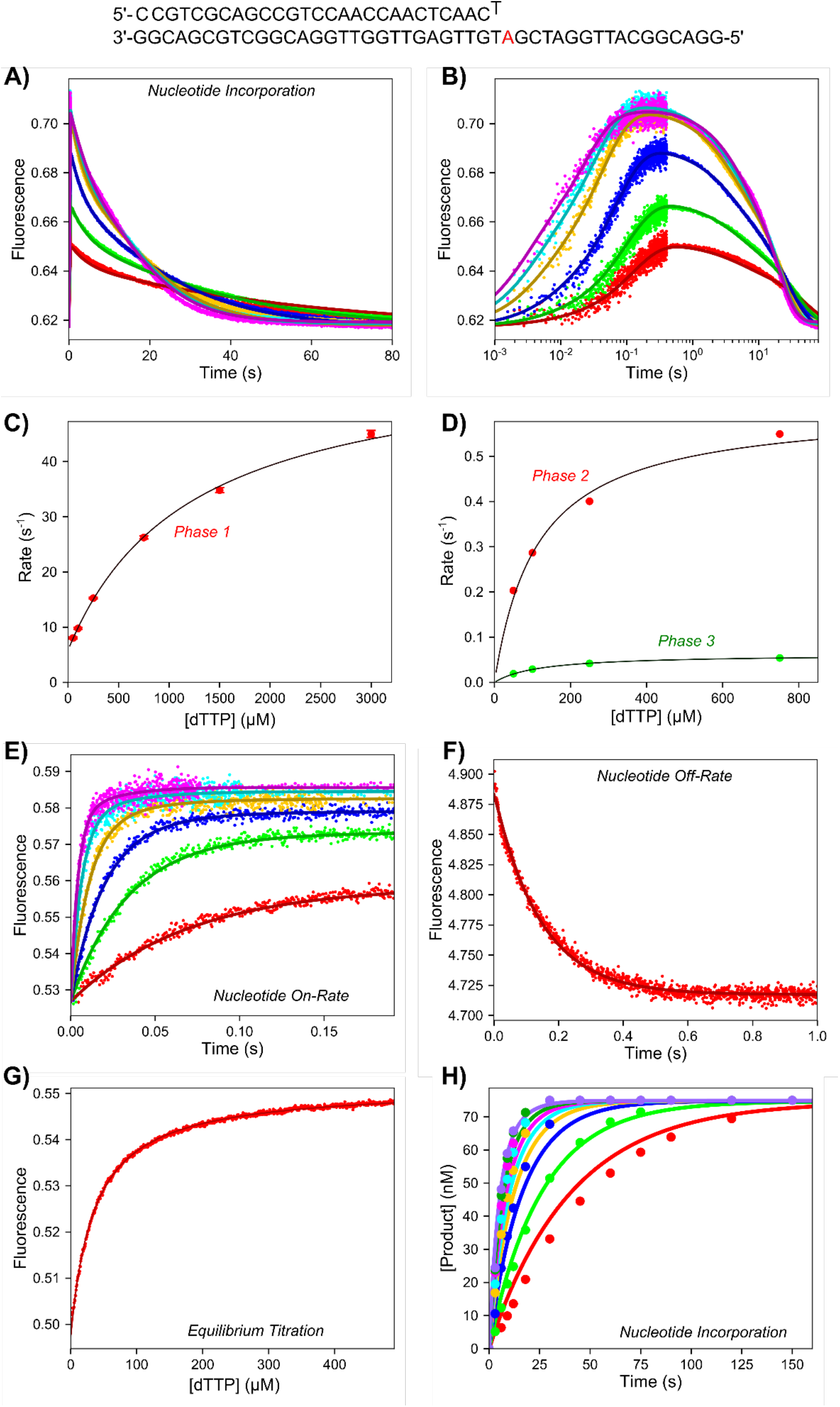
Conformational dynamics during the full dTTP mismatch extension reaction. <u>DNA Substrate:</u> The DNA substrate used in the experiment in (A) is shown. <u>A) Stopped-flow dTTP mismatch extension reaction.</u> A solution of 100 nM T7 DNA polymerase E514Cou, 2 μM thioredoxin, and 150 nM DNA 27-T/45-18T was mixed with 0.05–3 mM Mg^2+^-dTTP and 12.5 mM Mg^2+^ to start the reaction. The solid lines are the best global fit of the data including the experiments in Figure 3. <u>B) Stopped-</u> <u>flow dTTP mismatch extension reaction–log time scale.</u> Same data as in (A) but shown on a logarithmic timescale to better reveal the fast phase of the reaction. <u>C) Rate versus concentration for the fast phase of the mismatch extension reaction.</u> Rates are from the 3-exponential fit of the data in (A) (not shown in panel A). Fitting the data to a hyperbola gives the parameters: y-intercept = 5.85 ± 0.32 s^−1^, amplitude = 54 ± 2 s^−1^, *K_d,app_* = 1.21 ± 0.12 mM. <u>D) Rate versus concentration for the slower two</u> <u>phases of the mismatch extension reaction.</u> Rates are from the 3-exponential fit of the data in (A). At high concentrations there is an additional phase that complicates the fitting so rate versus concentration is only reported for concentrations up to 750 μM dTTP. Fitting the data to a hyperbola gives the parameters for the 2^nd^ phase (red data points): y-intercept = 0 s^−1^, amplitude = 0.61 ± 0.06 s^−1^, and Kd,app = 113 ± 30 μM. Fitting the data to a hyperbola fives the parameters for the 3rd phase (green data points): y-intercept = 0 s^−1^, amplitude = 0.062 ± 0.001 s^−1^, *K_d,app_* = 118 ± 7 μM. <u>E) Stopped-flow mismatch extension nucleotide binding-rate experiment.</u> Experimental conditions are given in Figure 3A. The solid lines through the data are the best global fit with the other experiments in this figure. <u>F) Stopped-flow mismatch extension nucleotide dissociation-rate experiment.</u> Experimental conditions are given in Figure 3C. The solid line through the data is the best global fit with the other experiments in this figure. <u>G) Mismatch extension equilibrium dTTP binding titration.</u> Experimental conditions are given in Figure 3D. The solid line through the data is the best global fit with the other experiments in this figure. <u>H) Chemical-quench dTTP mismatch extension experiment.</u> Experimental conditions are given in Figure 2C. The solid lines through the data are the best global fit with the other experiments in this figure using the model shown in Figure 5 with rate constants in Table 5.

We were able to obtain a reasonably good fit using the three-step binding model for the stopped-flow data alone, however this mechanism was not sufficient when including the chemical-quench data from Figure 2C. To fit the chemical-quench data along with the stopped-flow data, the model required an additional step where dTTP binds to the FDT state and activates the enzyme (forming the GDT’ state) to stimulate the rate of product formation (Figure 5). With this model, the chemical-quench and stopped-flow data for the mismatch extension reaction could be fit simultaneously along with the binding data (with DNA_dd_). To ensure consistency, we linked rate constants for ground state binding (ED ↔ EDT), the first conformational change (EDT ↔ FDT), and the activation step (FDT ↔ GDT’) so they were identical for equivalent reactions with DNA_dd_ and normal DNA. The remaining rate constants were treated as independent parameters. The activation step forming the GDT’ state accounts for the fourth phase observed at high nucleotide concentrations in the stopped-flow experiment in Figures 4A and Figure 4B. The global fit for all mismatch binding and extension experiments is given in Figure 4 (A, E, F, G, H), where the lines through the data give the best global fit using the model shown in Figure 5. The values for 1/*K_4_* ranged from 2.3—3 mM but was locked at a value of 2.6 mM to compute the confidence contour analysis for the remaining parameters, which were reasonably well constrained by the data (Figure 6) with the best fit values listed in Table 5.

**Figure 5:**
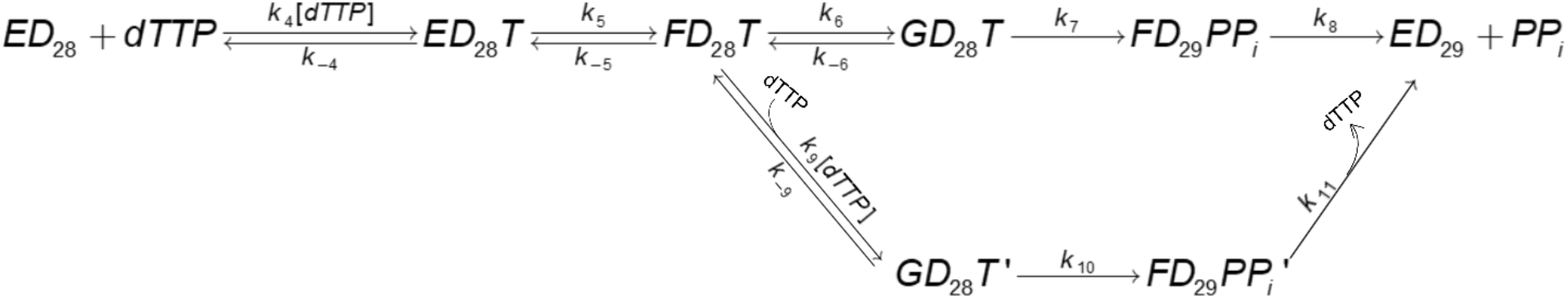
Kinetic scheme for dTTP mismatch extension reaction. The kinetic model used for data fitting for the experiments in Figure 4 is shown.

**Table 5:**
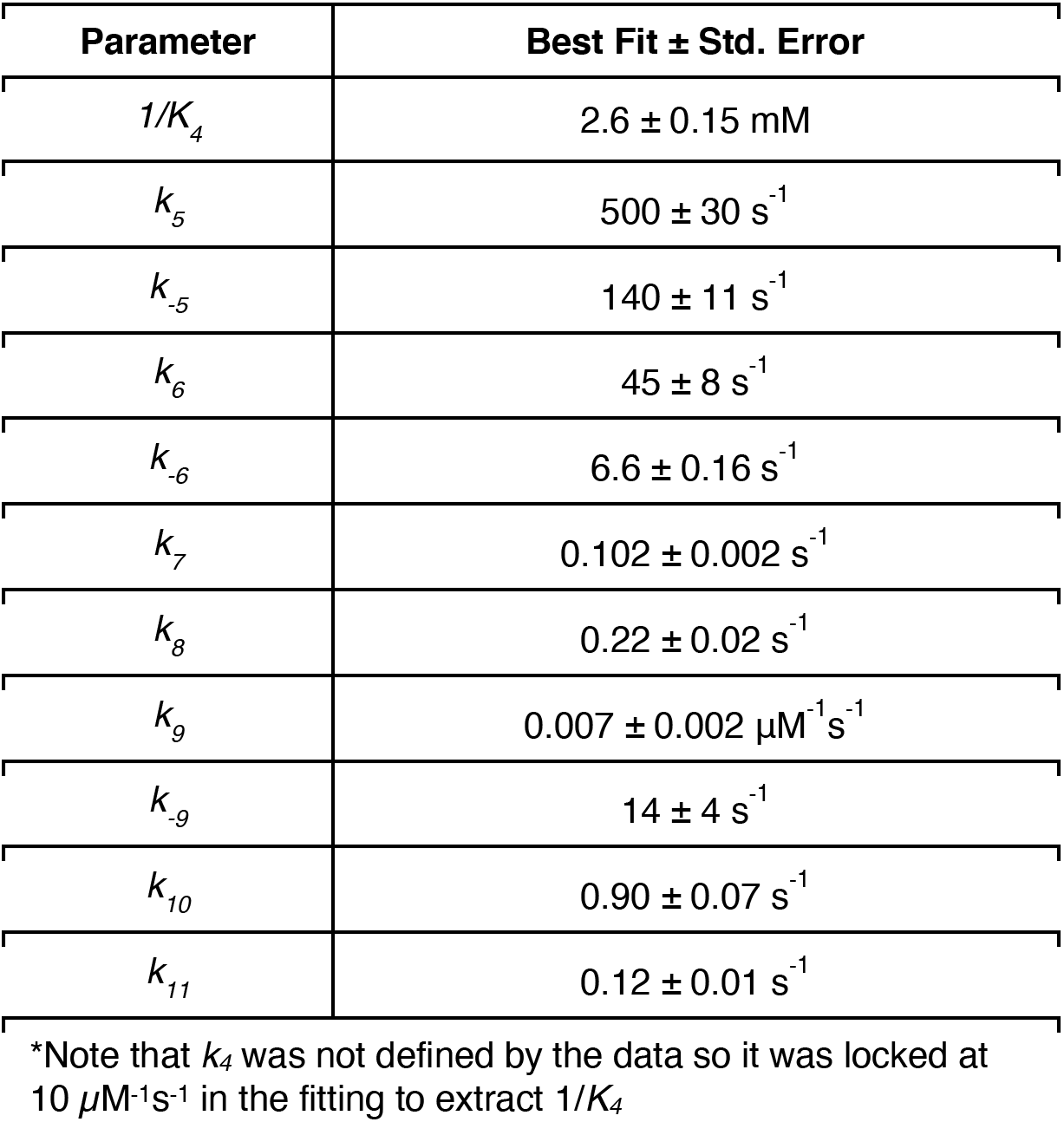
Parameters derived from global fitting mismatch extension data.

**Figure 6:**
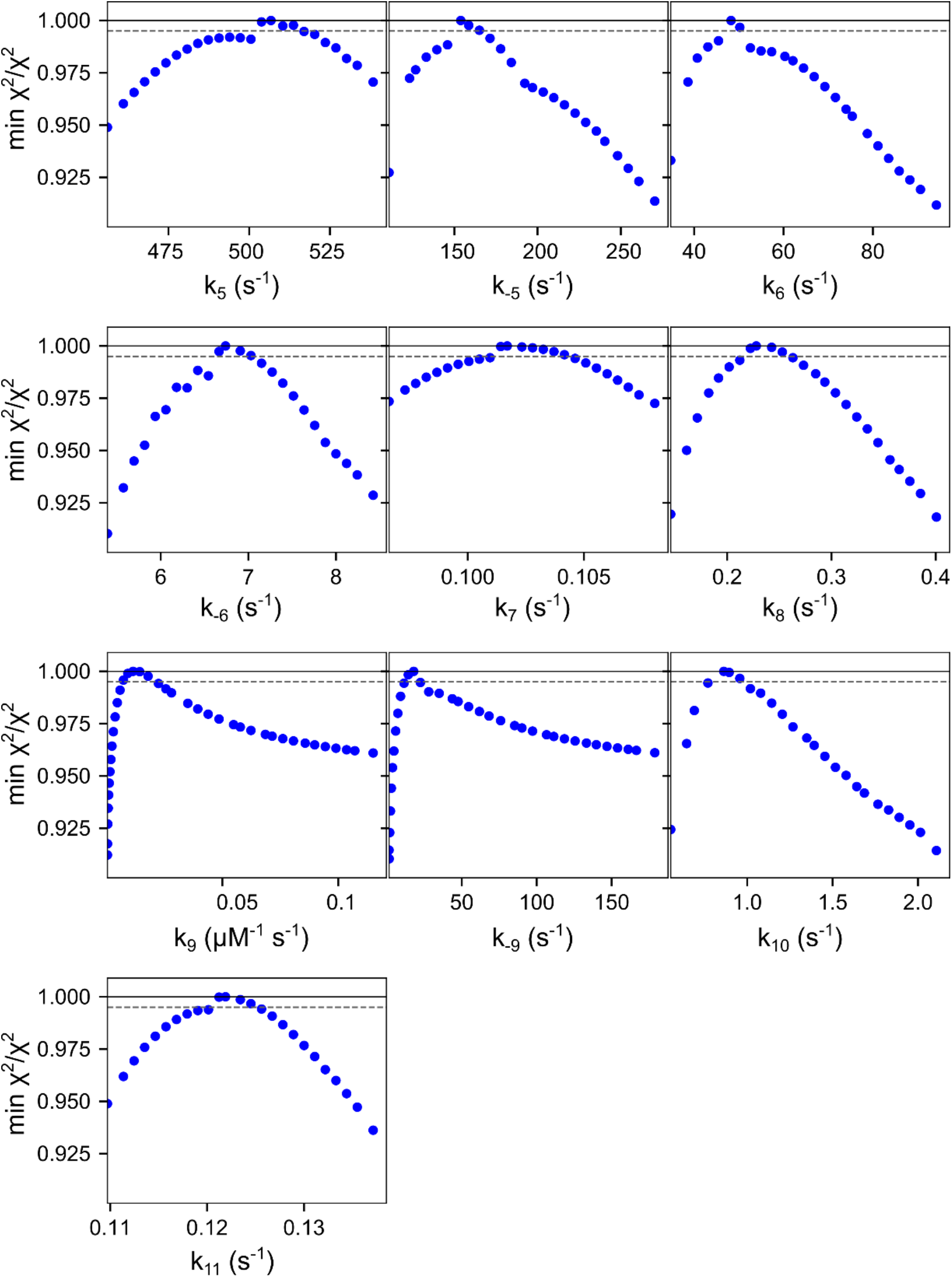
Mismatch extension confidence contours. Confidence contours from the FitSpace calculation in KinTek Explorer are shown for the global fit of the mismatch extension data in Figure 4. During initial exploration we found that the confidence interval for 1/*K_4_* ranged from 2.3 to 3 mM, but large variations in *K_4_* caused wider variations on other parameters. Therefore, we locked 1/*K_4_* = 2.6 mM to obtain the confidence contours shown here. The dashed grey line gives the *χ*^2^ threshold (0.995) reflecting the 95% confidence interval. Values from the confidence contour analysis can be found in Table 5.

### 2.5. Kinetics of dTTP:T misincorporation

With a kinetic model that describes mismatch extension, we proceeded to determine the kinetics of conformational changes occurring during the complete dTTP misincorporation and mismatch extension reaction. A stopped-flow experiment was performed by mixing a solution of T7 DNA polymerase E514Cou-DNA (27/45-18T, DNA in Figure 7) complex with 0.5–4 mM Mg^2+^dTTP and 12.5 mM Mg^2+^ (Figure 7A and 7B). The data show an initial decrease in fluorescence that can be attributed to the conformational changes occurring during misincorporation, followed by an increase in fluorescence at longer times and a return to the starting fluorescence level, consistent with the mismatch extension data. Conventional fitting of the data was challenging as the data fit 4 exponentials at low dTTP concentrations but at least 5 exponentials at high dTTP concentrations and there were large errors on the fitted parameters. For this reason, we skip showing the conventional fitting and fit the data directly by simulation using KinTek Explorer. Rate constants for the mismatch extension reaction were locked at best fit values determined from the global fitting of the data in Figure 4. Only rate constants for the misincorporation reaction were allowed to float as independent parameters during data fitting, and fitting was limiting to the initial phase where the fluorescence decreases then increases again.

**Figure 7:**
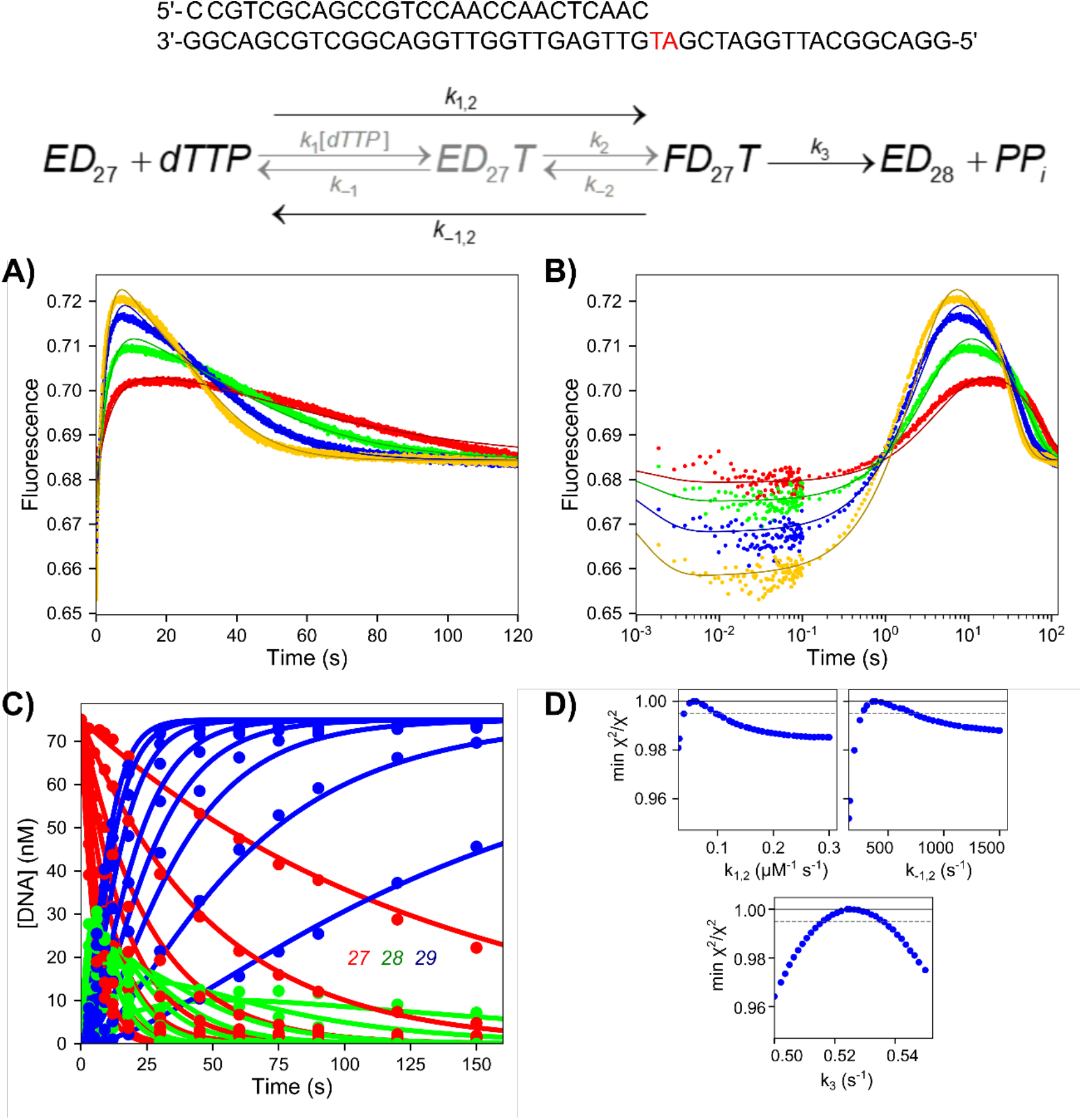
Conformational dynamics during misincorporation followed by mismatch extension. <u>DNA Substrate:</u> The DNA substrate used in the experiment in (A) is shown at the top. <u>Scheme:</u> Kinetic model for dTTP misincorporation. One step nucleotide binding (pathway in black) or estimated 2 step binding (pathway in grey) is followed by a single irreversible chemistry step. The mismatch extension scheme is not shown here but can be found in Figure 5. <u>A) Stopped-flow misincorporation and</u> <u>mismatch extension reaction.</u> A solution of 100 nM T7 DNA polymerase E514Cou, 2 μM thioredoxin, and 150 nM 27/45-18T DNA was mixed with 0.5–4 mM Mg^2+^-dTTP and 12.5 mM Mg^2+^ to start the reaction in the stopped flow. Solid lines through the data are the best fits by simulation, where rate constants from the misincorporation were allowed to float during data fitting, and rate constants for mismatch extension were locked at values derived from the data in Figure 4. <u>B) Stopped-flow misincorporation</u> <u>and mismatch extension reaction–log timescale.</u> Same data as in (A) but shown on a logarithmic time scale to better display the initial fast phase of the reaction, corresponding to the misincorporation step. <u>C) Chemical-quench dTTP misincorporation and</u> <u>mismatch extension reaction.</u> Reaction conditions are given in the figure legend from Figure 2A. The data are shown here, globally fit with the stopped-flow data, where the solid lines show the best fit by simulation. <u>D) Confidence contours for dTTP</u> <u>misincorporation reaction.</u> Experiments in this panel were fit to the scheme at the top of the figure to extract rate constants for the misincorporation reaction. Parameters for mismatch extension were determined in Figure 4 and are given in Table 5. The dashed grey line gives the *χ*^2^ threshold corresponding to the 95% confidence interval. The values from the confidence contour analysis can be found in Table 6.

The data fit the simple model shown in Figure 7 with one step nucleotide binding (*k_1,2_*) to form the FD27dTTP state, followed by a combined chemistry/PPi release step (*k_3_*). The stopped-flow data were globally fit with the chemical-quench data (Figure 7C) described in Figure 2A and confidence contour analysis was performed (Figure 7D) to determine whether the rate constants were well constrained by the data. As shown with a *χ*^2^min/*χ*^2^ threshold of 0.995 (to estimate the 95% confidence interval), each parameter was well constrained by the data. The apparent second order rate constant for nucleotide binding (*k_1,2_*) is relatively slow at 0.055 μM^−1^s^−1^, while the nucleotide dissociation rate (*k_−1,2_*, 340 s^−1^) is fast. Combining these rate constants provides an estimate of *K_m_* = 6.2 mM, consistent with the data presented in Figure 2. Parameters from the confidence contour analysis are given in Table 6. For comparison to rate constants derived for correct nucleotide incorporation with a 2-step nucleotide binding model, we estimated rate constants for a 2-step nucleotide binding model for the misincorporation reaction. We locked the nucleotide dissociation rate, *k_−2_*, at 340 s^−1^, and locked the second order rate constant for dNTP binding (*k_1_*) at a diffusion limited 100 μM^−1^ s^−1^. With a *K_m_* of 6.2 mM based on the chemical quench data in Figure 2, the fit by simulation showed ground state dNTP binding dissociation constant (1/*k_1_*) must be at least 9.3 mM so we locked *k_−1_* at 930,000 s^−1^ and fit to get the rate of the conformational change, *k_2_*, at 170 s^−1^. Estimated rate constants for the two-step nucleotide binding model are given in Table 6. Assuming rapid equilibrium nucleotide binding, *k_2_* is now smaller than *k_−2_*, (with very small *k_3_*) so the equilibrium constant favors nucleotide release and reduces the rate of incorporation by shifting the equilibrium away from the *FDN* state.

**Table 6:** Kinetic parameters for dTTP misincorporation.

| Model | Parameter | Best Fit ± Std. Error |
| --- | --- | --- |
| One-Step Nucleotide Binding | $k_{1,2}$ | $0.055 \pm 0.011 \mu\text{M}^{-1}\text{s}^{-1}$ |
| | $k_{-1,2}$ | $340 \pm 120 \text{ s}^{-1}$ |
| | $K_m$ | 6.2 mM |
| | $k_{cat}$ | $0.53 \pm 0.01 \text{ s}^{-1}$ |
| Two-Step Nucleotide Binding | $k_1$ | $100 \mu\text{M}^{-1}\text{s}^{-1}$ |
| | $k_{-1}$ | $930,000 \text{ s}^{-1}$ |
| | $1/K_1$ | 9.3 mM |
| | $k_2$ | $170 \text{ s}^{-1}$ |
| | $k_{-2}$ | $340 \text{ s}^{-1}$ |
| | $k_3$ | $1.6 \text{ s}^{-1}$ |

The $\chi^2$ threshold was set at 0.995 to get the 95% confidence interval for the one-step nucleotide binding model. Two step nucleotide binding parameters are based on estimates for ground-state binding dissociation constant ( $1/K_1$ ) of 9 mM and the $K_m$ of 6.2 mM from fitting the data using a one-step binding model.

According to these results we can see how the nucleotide-induced conformational change improves fidelity by slowing the rate constants leading to incorporation and favoring release of the bound mismatch. When *k_−2_* >> *k_3_*, the kinetic parameters for the three-step model can be approximated by the following terms:

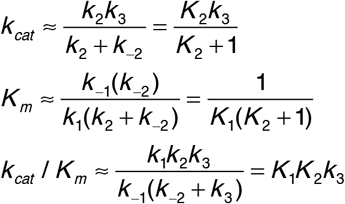

Accordingly, *k_cat_* is reduced by the fraction of enzyme in the FDN state, defined by *k_2_*/(*k_2_*+1), so with *k_2_* = 0.5 only one third of the bound nucleotide will be in the state to promote catalysis. The *K_m_* value is reduced relative to 1/*k_1_* by the term (*k_2_*+1). In this case, the *K_m_* = *K_d_* for nucleotide binding due to the rapid-equilibrium two-step binding. Finally, in *k_cat_/K_m_*, the (*k_2_*+1) term in both *k_cat_* and *K_m_* cancels, leaving the simple relationship, *k_cat_/K_m_* = *K_1_K_2_k_3_*. Therefore, values of *k_2_* < 1 reduce the specificity constant directly even though the *K_m_* is reduced by tighter binding. Note that *k_cat_* is not simply *k_3_* and *K_m_* is not 1/*K_1_K_2_*. The change in specificity-determining steps comparing correct versus mismatch base pairs is illustrated by considering the free energy profiles.

### 2.6. Free energy profiles comparing correct, mismatch, and mismatch extension reactions for T7 DNA polymerase

To compare the pathways for correct nucleotide incorporation, misincorporation, and mismatch extension, we constructed free energy profiles using the free energy profile tool in KinTek Explorer based on simple transition state theory. Rate constants determined by global fitting for each scenario were used in the calculation at a temperature of 293 K, and with a transmission coefficient of 0.01 to better visualize the profiles. Physiological concentrations of nucleotide were included in the calculation of the pseudo-first order rate constant for nucleotide binding (175 μM) based on previously estimates from *E. coli* (43). We also derived expressions for the parameters *k_cat_*, *K_m_*, and *k_cat_*/*K_m_* for each reaction scheme using the King-Altman method (44) and they are listed to the right of each free energy profile in Figure 8. For comparison we also present the free energy profile for correct nucleotide incorporation in Figure 8A from rate constants determined in our previous study on the T7 DNA polymerase E514Cou variant (31), without the translocation and competitive PPi binding for simplicity. Although we recognize the translocation step must be occurring in our model for misincorporation, the experimental data do not define this step based on the experiments presented in this paper and it is expected to be much faster than the misincorporation reaction. In the profile for correct incorporation, the highest barrier relative to the starting material (ED + A) is the barrier for the conformational change step (EDA to FDA), indicating that *k_cat_*/*K_m_* (the specificity constant) is determined by all steps leading up to and including the conformational change step. Simplifying the equation for *k_cat_*/*K_m_* based on the slow nucleotide dissociation rate (*k_−2_* ≪ *k_3_*), we show that *k_cat_*/*K_m_* ~ *K_1_K_2_*. Although the conformational change (*k_2_* = 6,500 s^−1^) is much faster than the rate of chemistry (*k_3_*, ~300 s^−1^), the nucleotide dissociation rate constant (*k_−2_*, 1.7 s^−1^) is much slower than the rate of chemistry, so the conformational change commits the correct nucleotide to incorporation so the conformational change step the highest energy barrier in the free energy profile.

**Figure 8:**
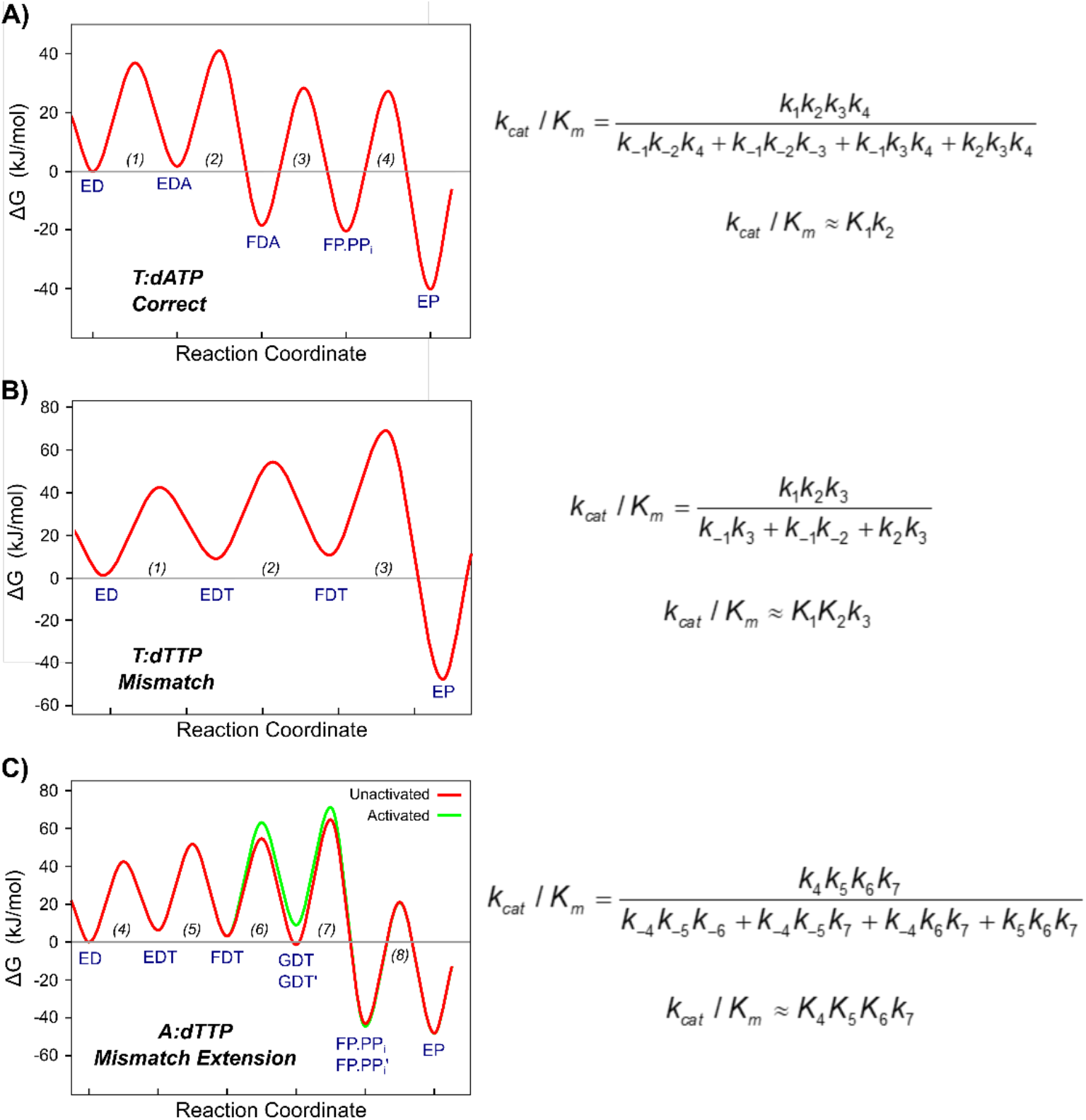
Free energy profiles comparing correct incorporation to misincorporation and mismatch extension. Free energy profiles were created in KinTek Explorer with the best fit rate constants derived from global fitting. Physiological nucleotide concentrations (175 μM dNTP (43)), a transmission coefficient = 0.01, and temperature = 293 K were used. Steady state kinetic parameters for each model are shown to the right of the free energy profiles and were determined with the King-Altman method. <u>A) Free energy profile for correct nucleotide incorporation.</u> Rate constants used to make this free energy profile are from (31). The highest barrier relative to the starting material is for the conformational change step (EDA to FDA states). <u>B)</u> <u>Free energy profile for dTTP misincorporation.</u> The highest barrier relative to the starting material is the chemistry step (FDT to EP state). <u>C) Free energy profile for mismatch extension.</u> The highest barrier relative to the starting material is chemistry (GDT to FP.PPi and GDT’ to FP.PPi’ states). The green curve is the free energy profile for the activated form of the enzyme (GDT’).

The free energy profile for the misincorporation reaction is shown in Figure 8B using the estimated parameters for a two-step nucleotide binding model for direct comparison to the other free energy profiles. The highest barrier relative to the starting material is the chemistry step (FDT to EP), indicating that specificity is determined by all steps leading up to and including the chemistry step. It is also important to note that the stopped-flow data for the misincorporation reaction showed a decrease in fluorescence, rather than an increase in fluorescence seen for the correct nucleotide. This indicates that the mismatched nucleotide may not go through the same conformational pathway as the correct nucleotide, but rather may proceed through a separate pathway to reach a different state. This result suggests that the enzyme enters a different state after binding a mismatch compared to the state formed after binding a correct nucleotide, as reported by the fluorescence signal. The plausible conclusion from this observation is that after binding a mismatch, the conformational change step serves to disorganize catalytic residues to reduce the rate of misincorporation. The data suggesting the “closed” state following nucleotide binding is different for correct versus mismatched nucleotides provide another way in which the conformational change is a major determinant of enzyme specificity. In addition to promoting release of a bound nucleotide by failing to bind a mismatch tightly, the altered conformation may also function to reduce the rate of misincorporation.

The free energy profile for the mismatch extension reaction is shown in Figure 8C. This reaction proceeds through at least three binding steps, with each peak higher in free energy than the previous. The highest barrier relative to the starting material is the chemistry step (GDT to FP.PPi), indicating that, like the mismatch, specificity is determined by all steps leading up to and including the chemistry step which can be seen in the expression for *k_cat_*/*K_m_*. This reaction also required an activation step where a second dTTP molecule acts as an activator for the chemistry step, although the free energy profile demonstrates that this “activation” pathway is actually higher in free energy than the “unactivated” pathway.

We performed flux calculations in KinTek Explorer to determine the fraction of the FDT state that proceeds via the activated pathway through GDT’ versus the un-activated pathway through GDT (See Figure S2). At low, physiological nucleotide concentrations (up to 250 μM), less than 10% goes through the activated pathway. At high nucleotide concentrations (3 mM), over 60% goes through the activated pathway. The role of this activation step and the location of the weak nucleotide binding site remain as open questions and may not be relevant under physiological nucleotide concentrations. However, by including this step during global data fitting we resolved the slower phase so that the faster, more physiologically relevant steps could be more accurately defined.

### 2.7. Computer simulation characterization of bound mismatches

To better understand the structural basis for the weaker binding and slower rates of chemistry for the mismatch incorporation and mismatch extension relative to the correct nucleotide, all atom MD simulations were performed on each of these complexes as described in Materials and Methods. Although MD simulations do not allow us to directly simulate rates of the chemical reaction, the structures sampled in MD simulations provide insights to indicate alterations of the alignment of substrates and catalytic residues that are likely to impact the rates of catalysis.

Conformations sampled in equilibrium is used to investigate the overall structure of the domains in different complexes. The stable states of the correct- and mismatched-nucleotide complexes are shown in Figure 9 for comparison. For clarity, the mismatch extension complex is omitted because of its similarity to the mismatched-nucleotide complex. The overall conformation of the DNA, the thioredoxin binding domain (TBD), thioredoxin, the thumb domain, and the 3’-5’ exonuclease domain did not show any major differences between the different complexes. In all complexes, the largest differences are found in the “fingers” domain of the protein in comparing the correct nucleotide state with the mismatch recognition state. This is the region where the fluorescent unnatural amino acid is located (at position 514) to monitor conformational changes in our experiments. The conformation of the fingers domain with the correct nucleotide bound stayed in a closed state (shown in green in Figure 9) during the simulation timeframe, with the fingers domain and catalytic residues wrapped tightly around the nucleotide substrate. On the other hand, the conformation of the “fingers” domain for the complex with the mismatched nucleotide adopt a more open conformation. Here, the catalytic residues positioned farther away from the bound nucleotide to accommodate the mismatch. The more open conformation and repositioning of the catalytic residues seen with the mismatch explains the 200-fold slower rates of incorporation of a mismatch compared to a correct nucleotide. The structural changes shown by the MD simulations also explain the different direction of the changes in fluorescence observed for a mismatch compared to a correct nucleotide. All together, simulations support our model in which the enzyme follows a different conformational pathway for the nucleotide binding and incorporation of a correct nucleotide versus a mismatch.

**Figure 9:**
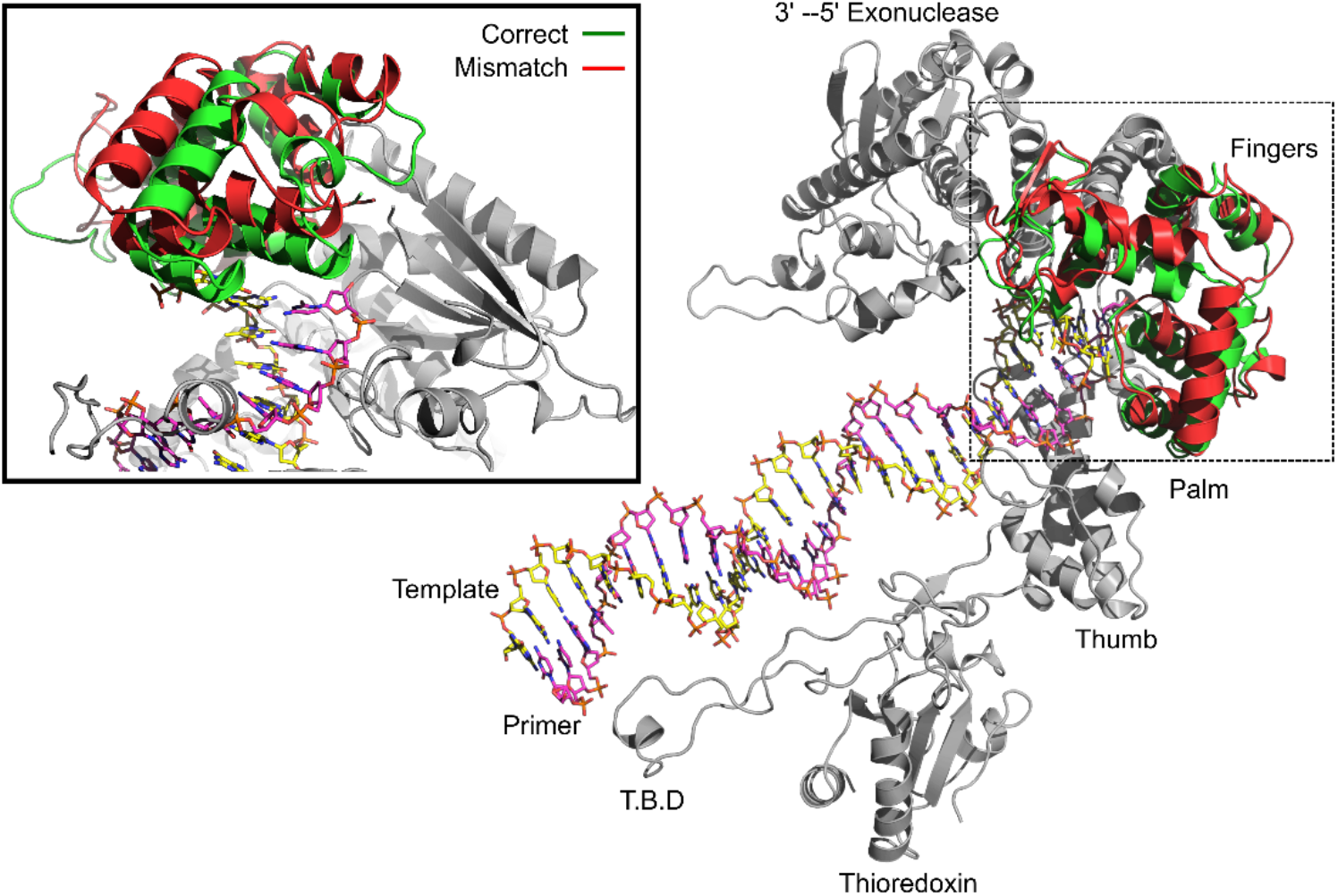
Conformations of correct nucleotide and mismatched nucleotide complexes. Conformations of enzyme-DNA-dNTP ternary complexes were determined by all atom molecular dynamics simulations with correct and mismatched nucleotides. The overall structure of the enzyme is given on the right, with domains of the enzyme labeled. Conformations of the correct- and mismatched-nucleotide complexes were similar in all regions of the protein except the fingers domain so these more stable regions are shown in grey. The fingers domain for the bound correct nucleotide complex (T:dATP) is shown in green and the fingers domain for the bound mismatched nucleotide complex (T:dTTP) is shown in red. A close-up view of the active site is shown in the inset on the left side of the figure. The mismatch complex displays a conformation more similar to the open conformation than the correct nucleotide. Since the overall conformation of the bound mismatch extension complex was almost identical to the mismatch complex, only the mismatch structure is shown in the figure for simplicity. Views of the active site are shown in Figure 10.

The stable structure seen for the binding of a correct nucleotide on top of a mismatch (leading to the mismatch extension reaction) shows parallels with the mismatch recognition state, adopting an open fingers conformation. It is still unclear why the mismatch extension shows an increase in fluorescence upon nucleotide binding if the overall conformation is similar to the mismatch complex, but it is known that even subtle differences in the residues surrounding the fluorophore can lead to these changes. As the enzymes closes around a correct nucleotide on top of a mismatch, it is reasonable to expect to see an altered structure that differs from either correct or mismatched nucleotides at the polymerase active site.

Closer examination of the base pairing of the incoming nucleotide in each complex, along with the alignment of nearby catalytic residues, provides a structural rationale for why mismatches are incorporated slower and bind with a lower affinity than correct nucleotides. Figure 10 provides a closer look at the active site, represented by average positioning of the incoming nucleotides. Figure 11 and Table 7 on the other hand provide quantitative assessment of several key properties for each complexes shown in Figure 10. In the correct nucleotide complex an average of 2 hydrogen bonds are observed during the entire 500 ns trajectory with the average angle of the hydrogen bonds around 5°. This geometry is close to perfect alignment and consistent with the 2 hydrogen bonds expected for an A:T base pair. In this state, the 3’-OH of the primer strand is 3.1 ± 0.04 Å away from the α-phosphate of the incoming nucleotide and is properly aligned for nucleophilic attack. In addition, the catalytic residues for this complex (Figure 10B) all make optimal contacts with the incoming nucleotide, including the catalytically important R518.

**Figure 10:**
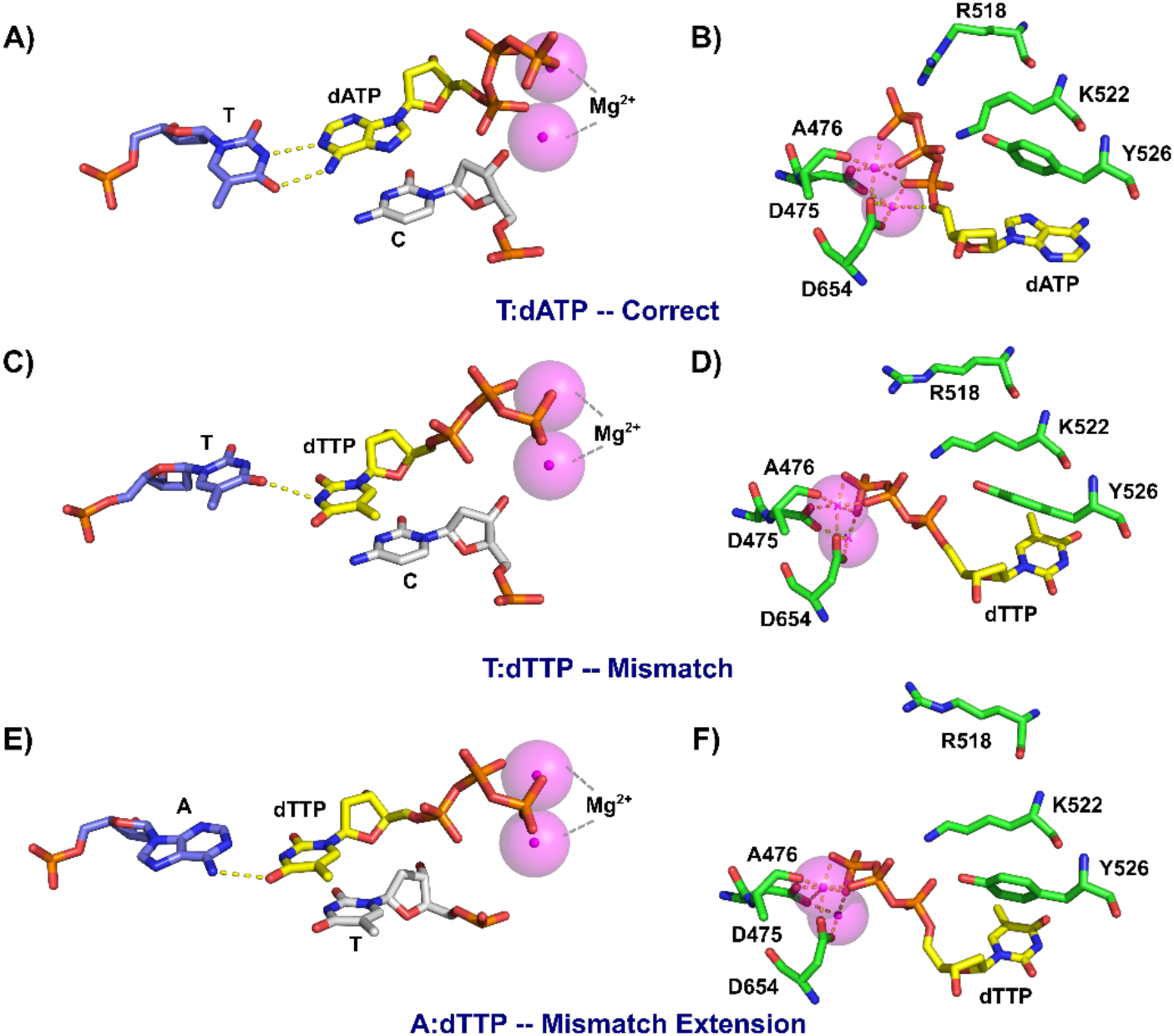
Base pairing and active site residue alignment of correct, mismatch, and mismatch extension complexes. Structures of each complex were determined by all atom MD simulations. <u>A) Base pairing of the correct nucleotide complex.</u> The incoming nucleotide is shown in yellow, the 3’ terminal base of the primer is shown in grey to demonstrate the alignment of the 3’-OH with the α-phosphate of the incoming nucleotide, the templating base is shown in blue, Mg^2+^ ions are shown in magenta, and hydrogen bonds are shown as dashed yellow lines. <u>B) Active site residue alignment of the correct nucleotide</u> <u>complex.</u> Protein residues are shown in green and are labeled. <u>C) Base pairing of the T:dTTP mismatch complex.</u> DNA bases are colored as in (A). <u>D) Active site residue alignment of the T:dTTP mismatch complex.</u> Colored as in (B). <u>E) Base pairing of</u> <u>the dTTP mismatch extension complex.</u> Colored as in (A). <u>F) Active site residue alignment of the dTTP mismatch extension</u> <u>complex.</u> Colored as in (B)

**Figure 11:**
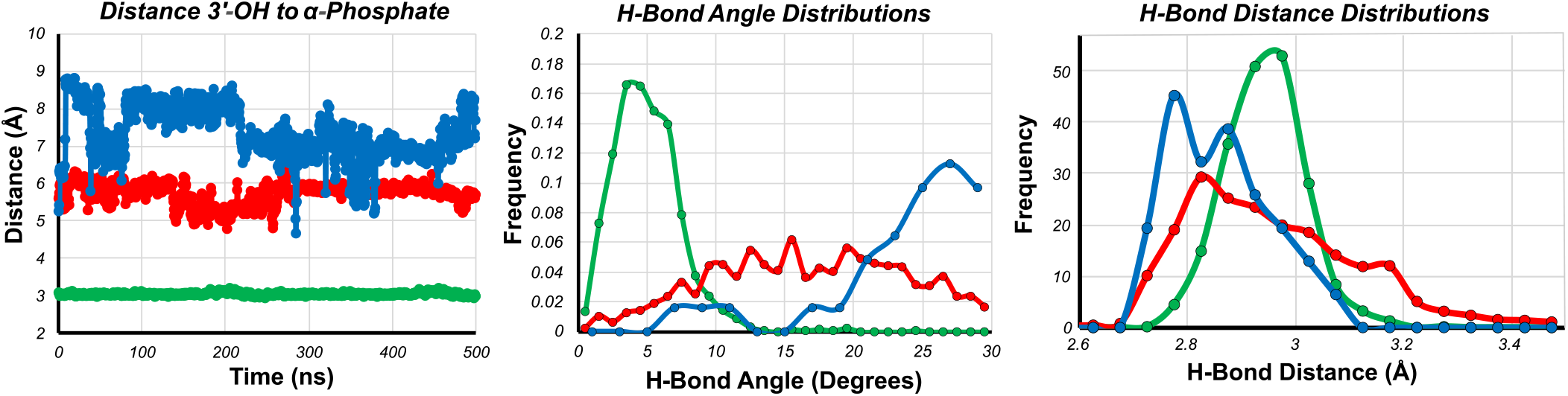
Analysis of the catalytic and h-bonding distances derived from MD simulations. Traces shown in green are for the *T:dATP-Correct* complex, red traces are for the *T:dTTP-Mismatch* complex, and blue traces are for the *T:dATP-Mismatch Extension* complex. The average of the parameters displayed here are summarized in Table 7. <u>A) Time evolution of the distance</u> <u>of 3’-OH to α-phosphate.</u> The distance from the 3’-OH of the primer to the α-phosphate of the incoming dNTP is shown as a function of time for each complex. <u>B) Hydrogen bond angle distribution.</u> The distribution of hydrogen bond angles between the incoming dNTP and the templating base are shown for each complex. <u>C) Hydrogen bond distance distributions.</u> The distribution of hydrogen bond distances between the incoming dNTP and the templating base are shown for each complex.

**Table 7:** Average parameters from quantitative analysis of MD simulation trajectories.

| Complex | Distance 3'-OH to α-Phosphate (Å) | Hydrogen Bond Angle (°) | Hydrogen Bond Distance (Å) | # of Hydrogen Bonds |
| --- | --- | --- | --- | --- |
| T:dATP (Correct) | $3.05 \pm 0.04$ | $5.05 \pm 2.11$ | $2.94 \pm 1.54$ | $2 \pm 0$ |
| T:dTTP (Mismatch) | $5.73 \pm 0.29$ | $16.62 \pm 4.00$ | $2.96 \pm 1.64$ | $1.01 \pm 0.71$ |
| A:dTTP (Mismatch Extension) | $7.36 \pm 0.69$ | $23.73 \pm 4.56$ | $2.86 \pm 1.56$ | $0.025 \pm 0.156$ |

Base pairing and primer alignment for the T:dTTP mismatch complex are shown in Figure 10C for comparison. Here, the dTTP with the templating T is impaired, resulted in an average of 1 ± 0.7 hydrogen bonds and with an average hydrogen bond angle of 17 ± 4° (Table 7). The 3’-OH group of the primer strand is also misaligned, with an average distance from the α-phosphate of the incoming nucleotide of 5.7 ± 0.3 Å. The fluctuation on these parameters are also larger than the fluctuations for the correct nucleotide complex and reflect the destabilization and disorder of the bound mismatch (full time course data are shown in Figure 11). The catalytic residues show an increase in distance for the mismatch complex (Figure 10D). Small increases in distances of reactive groups observed in the MD simulations are sufficient to explain the slowed rate of misincorporation. The phosphates of the incoming nucleotide are in a different conformation than seen for the correct nucleotide complex, hindering alignment with the Mg^2+^ ions coordinated to D475, A476, and D654. Interestingly, R518 is severely misaligned and does not form proper contacts with the incoming nucleotide. The Y526 residue is also misaligned to compensate for the conformation of the nucleotide.

The bound dTTP for the mismatch extension is shown in Figure 10E. The altered base pairing as well as increased distance of the 3’-OH of the primer to the α-phosphate of the incoming nucleotide is common to the mismatch complex. Although dTTP is the correct base for a templating A, the T:dTTP mismatch at the primer terminus distorts alignment of the bases in the active site with an average hydrogen bond angle of 24 ± 4.6°. In this state the hydrogen bonding stayed in an average of 0.02 ± 0.16 during the simulation (Figure 11, Table 7). The 3’-OH is also severely misaligned, with an average distance of 7.4 ± 0.7 Å that is farther away from a proper attack. This result is consistent with the observed reaction being even slower than the misincorporation reaction. The alignment of catalytic residues in Figure 10F shows that like the mismatch, R518 is significantly misaligned; however, the alignment of Y526 is better than for the mismatched-nucleotide complex. While mismatched nucleotide binding is weakened, larger effects can be seen on the alignment of catalytic residues and alignment of the 3’-OH, which is known to contribute to the greatly reduced rate of chemistry.

## 3. Discussion

This study completes our prior characterization of the role of enzyme conformational dynamics in DNA polymerase nucleotide specificity (12,13,31,42) by extending our analysis of a high fidelity DNA polymerase to include the kinetics of misincorporation. Prior analysis of nucleotide-induced conformational changes using the T7 DNA polymerase E514Cou variant showed that for a correct base pair, the transition from the open to closed enzyme state was at least 20-fold faster than chemistry, but the reverse conformational change to allow release of the nucleotide was much slower than chemistry. Accordingly, the conformational change commits a correct nucleotide to incorporation so that the specificity constant for correct nucleotide incorporation is defined solely by the equilibrium constant for initial nucleotide binding and the rate constant for the conformational change step (*k_cat_/K_m_* = *K_1_K_2_*) (30). DNA polymerases, however, must not only incorporate correct nucleotides efficiently, but they must also effectively discriminate against mismatched nucleotides and change specificity depending on the template. Here, we address the question of the role of conformational changes in discrimination against mismatch incorporation using the T7 DNA polymerase E514Cou variant to provide a fluorescence signal to measure conformational dynamics. Our data show that for misincorporation the rate of the chemistry step is significantly slower while the rate constant for nucleotide release is much faster than observed for a correct base pair. Therefore, the specificity constant is a function of the product of the equilibrium constants for binding and isomerization and the rate constant for the chemistry step (*k_cat_/K_m_* = *K_1_K_2_k_3_*). Accordingly, the discrimination index, defined by a ratio *k_cat_/K_m_* values, is a function different reaction steps for correct versus mismatched nucleotides.

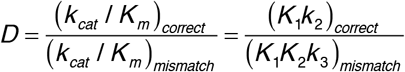

For a mismatched nucleotide, the unfavorable equilibrium constant for the conformational change step (*k_2_*) reduces *k_cat_* and *k_cat_/K_m_* by shifting the internal equilibrium away from the catalytic state while promoting nucleotide release.

In this paper we compared the free energy profiles observed for mismatch incorporation with the pathway for correct nucleotide incorporation derived previously (Figure 8). In the absence of this data, theoretical papers have used free energy profiles to argue whether or not induced-fit can contribute to specificity, leading to conflicting conclusions depending on the assumptions inherent in the analysis (11,45). Initial ideas for understanding polymerase fidelity were based on considering contributions due to changes in *k_cat_* and *K_m_* with the underlying—often unstated— premise that these parameters reflected changes in rates of incorporation and nucleotide affinity, respectively (46). However, our results demonstrate that the *K_m_* is a function of the nucleotide binding affinity only for mismatched, not for correct nucleotides. For correct nucleotide incorporation, *K_m_* = *k_3_*/*K_1_K_2_*, while nucleotide binding affinity (if the two-step binding reaction came to equilibrium) is defined by *K_d,net_* = 1/(*k_1_*(1+*k_2_*)). In contrast, for mismatched nucleotides, *K_m_* = *K_d,net_* = 1/(*k_1_*(1+*k_2_*)) because chemistry is slow allowing the conformational change to come to equilibrium.

A popular theoretical construct invoked checkpoints so fidelity was the product of contributions from each point along the pathway; namely, nucleotide binding, conformational changes, and chemistry (47). Although it was noted that the contribution of each step to overall selectivity can vary for different polymerases, we show that the contributions of each step to *k_cat_*/*K_m_* is different for correct and incorrect base pairs, so overall selectivity is not a simply a product of the selectivity at each individual step. Similarly, Fersht argued that a two-step binding cannot contribute more to fidelity than a thermodynamically-equivalent one-step binding (11), but this analysis assumes both binding steps are in rapid equilibrium, which is true for a mismatch (*k_−2_* ≫ *k_3_*) but is not true for correct incorporation where *k_−2_* ≪ *k_3_*. Arguments put forth by Warshel stating that a pre-chemistry conformational change cannot contribute to fidelity unless it is rate limiting (45) are incorrect because he confused rate-determining steps (*k_cat_*) with specificity-determining steps (*k_cat_*/*K_m_*). For correct incorporation, chemistry is rate limiting (*k_cat_* = *k_3_*), but the binding steps define specificity, *k_cat/Km_* = *K_1_k_2_*. One might ask why *k_cat_* is not included in the equation for *k_cat_*/*K_m_*. The answer is that when *k_−2_* ≪ *k_3_*, *k_3_* appears in both the numerator and denominator so it cancels in the equation for *k_cat_/K_m_* (48).

We can finally address the question of the role of induced-fit in specificity based on direct measurements as illustrated in Figure 8. For a correct nucleotide, the conformational change is fast (6500 s^−1^), and the nucleotide dissociation rate (1.7 s^−1^) is much slower than chemistry (~300 s^−1^). The slow dissociation rate brings the free energy well for the FDA state far down, committing the correct nucleotide to incorporation and so *k_cat_/K_m_* ≍ *K_1_K_2_*. For the mismatch (Figure 8B), the highest energy barrier is the chemistry step (between the FDT and EP states) so *k_cat_*/*K_m_* is determined by the product of the binding and chemistry steps and for the approximated 2 step model, *k_cat_*/*K_m_* ≈ *K_1_K_2_k_3_*. For the mismatch extension (Figure 8C), the highest barrier is also chemistry (between the GDT and FP.PP) states so *k_cat_*/*K_m_* for the three step nucleotide binding model is *k_cat_*/*K_m_* ≈ *K_1_K_2_K_3_k_4_*. In this case, both the conformational change and chemistry steps are crucial to specificity and appear in the expression for *k_cat_*/*K_m_*. In contrast, for correct nucleotide incorporation the rate of the chemical reaction is not included in the calculation of the specificity constant as long the chemistry is faster than the rate of enzyme opening to release the nucleotide.

The nucleotide-induced change in enzyme structure is the major determinant of enzyme specificity. The enzyme recognizes a correct nucleotide through a network of electrostatic interactions, binds the substrate tightly and organized catalytic residues to promote catalysis. In contrast, the enzyme fails to close tightly over a mismatched nucleotide and promotes enzyme opening and nucleotide release rather than catalysis. Specificity for moderate to high fidelity polymerases is largely attributable to the conformational change step to either bind a correct nucleotide tightly, organize catalytic residues, and commit the nucleotide to incorporation, or alternatively to bind a mismatch weakly, disorganize (or fail to organize) catalytic residues, and favor nucleotide release. Thus, kinetic partitioning of the FDN state defines specificity by either favoring incorporation of a correct nucleotide or release of a mismatch. This does not appear to be the case for low fidelity DNA polymerases such as Pol β where nucleotide binding, including the conformational change step, comes to equilibrium preceding incorporation of both correct and mismatched nucleotides (15,49). For high fidelity polymerases, the fast conformational change greatly increases fidelity and speed of incorporation.

Interestingly, the fluorescence signal during the misincorporation reaction decreases, whereas the fluorescence increases during the correct incorporation and mismatch extension reactions. This suggests a different conformational state for the bound mismatched nucleotide which is consistent with previous work on this enzyme and other labeled enzymes (48,50). The intriguing proposal arising from this work is that high fidelity enzymes may recognize a mismatch and use binding energy to misalign catalytic residues. Thus, a further enhancement of fidelity could be achieved more than from simply failing to tightly bind a mismatch. Our fluorescent label may be positioned to detect subtle changes in enzyme structure after binding a mismatched nucleotide which can change the FRET efficiency from nearby tryptophan residues to provide fluorescence signals in the opposite direction for correct versus mismatched nucleotide. For Pol β and *E. coli* DNA polymerase I Klenow fragment, various stopped-flow studies have been performed for mismatch incorporation where the signal change is in the same direction as for the correct nucleotide (15–18), but these are lower fidelity repair enzymes so smaller and/or different perturbations in enzyme structure during incorporation mismatches may be expected.

We screened all mismatch combinations to determine the range of discrimination values against various mismatches by T7 DNA polymerase. One might anticipate that the purine:purine mismatches would be least efficiently incorporated due to the two bulky bases that would seemingly sterically clash in the enzyme’s active site. However, the C:dCTP misincorporation was the least efficient with *k_cat_*/*K_m_* of 1.4 M^−1^s^−1^, 40–100 fold less efficient than the A:dATP and G:dGTP misincorporations. Structural studies on other polymerases show that this mismatch causes fraying of the primer/template junction and the two small bases are not close enough to form critical contacts (51,52). We also found that the T:dTTP incorporation was much more efficient than the C:dCTP misincorporation, with *k_cat_*/*K_m_* of 130 M^−1^s^−1^, despite the fact that this is also a pyrimidine:pyrimidine mismatch. We found that the mismatch with the lowest discrimination is the T:dGTP mismatch. This is consistent with previous studies measuring thermodynamic parameters for double strand formation that found among the most stable mismatches was the G:T mismatch, and among the least stable mismatches was the C:C mismatch (53). Wobble base pairing of the G:T mismatch, along with rare tautomeric forms of these bases have been proposed to explain the stability of this mismatch (52,53).

We examined the T:dTTP misincorporation reaction in more detail since it provided a usable fluorescence signal in the stopped-flow instrument. Since the templating base following misincorporation is an A, the T:dTTP mismatch is further extended to the 29 nt product before the enzyme stalls. While misincorporation reactions have been characterized in some detail, there is much less information about mismatch extension available in the literature, so this mismatch provided a unique opportunity to study both misincorporation and mismatch extension in a single reaction. We found that the mismatch extension reaction occurred with a *K_m_* approximately 10-fold lower than the *K_m_* for T:dTTP misincorporation. The value of *k_cat_* for the mismatch extension reaction, however, was approximately 3-fold lower than *k_cat_* for the misincorporation reaction. This is consistent with previous studies on this enzyme showing that extension of a mismatch with dCTP occurred with a lower *k_cat_* and lower *K_m_* than observed for misincorporation. We also found that the *K_m_* for mismatch extension was higher than for a correct base but much lower than for a mismatch, while *k_cat_* was greatly reduced relative to the rates of misincorporation (9).

To better understand the structural elements driving specificity, we performed all atom MD simulations in different nucleotide-bound states of the complex including the correct nucleotide complex, mismatch complex, and mismatch extension complex. So far, it has been difficult to obtain a crystal structure of any high fidelity DNA polymerase with a bound mismatch without using Mn^2+^ (52,54), or mutagenesis to lower the fidelity of the enzyme (51). MD simulations have become much more accurate in the past decade and now provide an excellent method to not only study complexes with alternative substrates, but also to understand the dynamic motions of enzymes in different states (50,55). First, we investigated the differences in overall conformation between the different bound nucleotide states for T7 DNA polymerase. Compared to the correct-nucleotide state, the mismatched-nucleotide complex showed a much more open configuration, especially localized in the fingers domain. This is also consistent with our stopped-flow data, where the mismatched nucleotide induces a fluorescence change in a different direction than observed for the correct nucleotide. Closer examination of the base pairing geometry and catalytic residues near the active sites provided more insights into the induced-fit mechanism. The correct base was stably bound, with the expected 2 hydrogen bonds for an A:T base pair, and the MD simulation trajectory showed little movement of the base during the 500 ns time frame, reflecting the proper alignment of catalytic residues with the incoming nucleotide. In contrast, the mismatched-nucleotide structure showed much more dynamic and significant movement of the nucleotide during the simulation. Other structural studies on mismatches have shown that the structure of each mismatch is distinct, and that some mismatches adopt conformations previously seen with DNA only, but some adopt new conformations (52). Another MD simulation study looked at the structural impact of DNA mismatches and showed that mismatches produced significant local structural alterations, especially in the case of purine transversions (56). They showed that mismatched base pairs often show promiscuous hydrogen bonding patterns which interchange among each other in the nanosecond timescale, consistent with the behavior we see in our trajectories. Furthermore, they showed that there is not a single path of mismatch-induced changes since different types of mismatches modify DNA properties in different ways. This is also consistent with our results since there are a wide range of rates of misincorporation and the mechanisms for mismatch discrimination vary between misincorporation and mismatch extension complexes. Of particular importance is not only hydrogen bonding, but more importantly how hydrogen bonding influences the alignment of the 3’-OH group with the α-phosphate of the incoming nucleotide. For both the mismatch and mismatch extension complexes, the alignment of the 3’-OH is severely perturbed to satisfy other hydrogen bonding within the active site. This has also been shown for other enzymes (54) and is likely a large contributor to the large decrease in the rate of mismatch incorporation.

The importance of base pair hydrogen bonding to confer optimal alignment for catalysis was questioned in earlier studies using nucleotide analogs that lacked hydrogen bonding capabilities and led to the conclusion that selectivity only depended on steric parameters to define a canonical base pair (57). However, more accurate kinetic analysis of the incorporation of these analogs demonstrated that hydrogen bonds contributed 2—7 kcal/mol toward nucleotide incorporation specificity and efficiency (58,59). The MD simulations presented here support the conclusion that hydrogen bonds provide an essential contribution to fidelity by enforcing correct base-pair geometry and alignment for catalysis.

For the past two decades, researchers in the DNA polymerase field have focused on whether the conformational change or chemistry was rate-limiting (45,47,60–64), which is the wrong question. The more important question is whether the reversal of the conformational change to allow release of bound nucleotide is slower than chemistry. As demonstrated by the free energy profiles, *k_cat_*/*K_m_* is determined by the rate constants for all steps leading up to and including the first largely irreversible step. For a correct nucleotide, the fast conformational change and a slow nucleotide off-rate essentially makes the conformational change step irreversible so *K_1_K_2_* defines specificity. For the mismatch and mismatch extension, chemistry is the first irreversible step in the pathway, so specificity is a function of all steps leading up to and including chemistry, as seen in the expression for *k_cat_*/*K_m_* for this pathway in Figure 8. Although we have not shown data for the reverse pyrophosphorolysis reaction here for a mismatch or mismatch extension, preliminary experiments suggest that pyrophosphorolysis is extremely slow and can be attributed to misalignment of the 3’ base of the primer which does not allow binding of PPi to the enzyme. Similarly, in our previous paper, we examined the forward reaction in the presence of PPi and saw significant inhibition. We proposed that PPi release occurs before translocation which has an equilibrium favoring the post-translocation state. When similar experiments were performed for a mismatch, there were no significant changes in the stopped-flow traces, suggesting the release of PPi is essentially irreversible.

We have shown the structural and kinetic bases for DNA polymerase fidelity in this paper. While we only investigated a single site for misincorporation reaction in detail, the underlying general principles derived in studies of this mismatch are likely to hold for most sequence contexts. Our labeling methods could likely be applied to other high-fidelity polymerases to test if the same fidelity mechanisms apply. The other contributing factor to fidelity is the 3’— 5’ proofreading exonuclease function. Ongoing investigations of the proofreading exonuclease function of this enzyme, combined with these data will give a more complete model of specificity for a high-fidelity DNA polymerase.

## 4. Materials and methods

### 4.1. Preparation of proteins/reagents

All experiments were performed with exonuclease deficient variants of T7 DNA polymerase (D5A/E7A) (29). Thioredoxin, wild-type T7 DNA polymerase, and T7 DNA polymerase E514Cou were expressed in *E. coli* and purified as previously described (30). BSA was purchased from New England Biolabs. Unless otherwise noted, a 20-fold molar excess of thioredoxin over T7 DNA polymerase was included. dNTPs were purchased from New England Biolabs. DTT was purchased from Gold Biosciences. All other buffer components were purchased from Fisher Scientific.

### 4.2. Preparation of oligonucleotides

All synthetic oligonucleotides were synthesized by Integrated DNA Technologies with standard desalting and were further purified in house by denaturing PAGE to a final purity of >99% full length oligo. Purified oligos were stored in 66.2 Buffer (6 mM Tris-HCl pH 7.5, 6 mM NaCl, 0.2 mM EDTA) at −20°C. Concentrations of purified oligos were determined by absorbance at 260 nm using the extinction coefficients given in Table 1. Double stranded DNA substrates for kinetic assays were prepared by mixing the appropriate primer with the appropriate template at a 1:1.05 molar ratio in Annealing Buffer (10 mM Tris-HCl pH 7.5, 50 mM NaCl, 1 mM EDTA), heating to 95°C, and slowly cooling to room temperature over 2 hours. Reactions with 2’,3’ dideoxy terminated primers annealed to 45 nt template (45-18N in Table 1, where N represents the templating base), designated DNA_dd_, were used to measure nucleotide binding without the chemistry step.

### 4.3. Reaction conditions

All experiments were carried out in T7 Reaction Buffer (29) (40 mM Tris-HCl, pH 7.5, 50 mM NaCl, 1 mM DTT, 1 mM EDTA). All reactions were performed with 12.5 mM Mg^2+^ during the reaction. Unless otherwise noted, reactions were set up by preincubating enzyme with DNA in the absence of MgCl_2_ and mixing with a solution of dNTP and MgCl_2_ to start the reaction. Nucleotide solutions were complexed with magnesium to form Mg2+-dNTP’s before use in kinetics experiments with high nucleotide concentrations. Reaction components given in the text are concentrations after mixing. All experiments were performed at 20°C.

#### 4.3.1. Chemical-quench experiments

Chemical-quench experiments were performed by hand mixing a solution containing the enzyme/[6-FAM]-DNA complex with a solution containing nucleotide and Mg^2+^ to start the reaction. Reactions were quenched by addition of EDTA to a final concentration of 0.3 M. To maintain a constant temperature of 20°C during the reaction, a modified tube rack designed to sit in a refrigerated water bath was used. Quenched samples were analyzed by capillary electrophoresis on an Applied Biosystems 3130xl Genetic Analyzer instrument with a 36 cm capillary array filled with POP-6 polymer (MCLab). Before injection, 1 μl of sample was mixed with 10 μl of HiDi Formamide (ThermoFisher) containing a Cy3 labeled oligo (Table 1) as an internal size standard. The fluorescence intensity of the FAM label was used to quantify the reaction products by peak integration with GeneMapper software (ThermoFisher) using previously reported methods (65).

#### 4.3.2. Stopped-flow experiments

Stopped-flow experiments were performed on a SF-300X instrument (KinTek Corporation, www.kintekcorp.com) with a circulating water bath for temperature control, a 150-watt xenon lamp as the light source, and a dead time of 1.3 milliseconds. Stopped-flow traces in the main text are averages of at least 6 individual traces and were repeated at least twice to ensure reproducibility. Fluorescence experiments with T7 DNA polymerase E514Cou were performed with excitation at 295 nm and emission at 445 nm, observed with a 45 nm bandpass filter (Semrock). Because of the excitation wavelength (295 nm) and high nucleotide concentrations that have absorbance at this wavelength, the inner filter effect became a significant problem. The *concentration series scaling* feature of KinTek Explorer was used to correct individual traces by adding a separate multiplier scaling factor for each trace at each concentration. For longer timescale experiments with a fast initial phase followed by a much slower phase, data were collected on two timescales to better resolve both reactions.

#### 4.3.3. Equilibrium titration measurements

Equilibrium titrations were performed with the TMX titration module for the SF-300X instrument with a solution of 280 μl of T7 DNA polymerase E514Cou, thioredoxin, DNA_dd_ and Mg2+ in the cuvette in a temperature-controlled chamber, with constant mixing with a micro stir bar. From a Hamilton syringe, 20.5 μl of titrant (7.5 mM Mg^2+^-dTTP) was added to the cuvette with constant mixing over the course of 5 minutes. The data were corrected for the small dilution before further processing. This dilution corrected data showed a decrease in fluorescence intensity at high dTTP concentrations following the initial increase, due to the inner filter effect. The data were therefore fitted in KinTek Explorer (for a simple 1 step binding model) with the observable A1*(E+ED+(b1*EDT))*(1-10^(-q*[dTTP]))/(2.303*q*[dTTP]), where *A1* is the scaling factor for the initial fluorescence, *b1* is the fluorescence scaling factor for the EDT state, and *q* is l/2*ɛ, where *l* is the path length and ɛ is the extinction coefficient. *q* was obtained from the initial fitting in KinTek Explorer and then used to correct the data for the inner filter effect. The corrected titration curve shown in Figure 3D was then fit using the observable A1*(E+ED+(b1*EDT)), where *A1* is the scaling factor for the starting fluorescence and *b1* is the fluorescence scaling factor for going to the EDT state. Titration data were then fit to the models described in the main text after correction.

### 4.4. Data fitting

Data fitting and analysis were performed with the simulation software KinTek Explorer v10 (35) (www.kintekexplorer.com). This software was also used in preparing figures for kinetic data. For conventional fitting to equations, the analytical fit *(aFit)* function of KinTek explorer was used. The hyperbolic equation used was 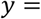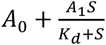, where *A0* is the y intercept, *A1* is the maximum value for the y axis variable, and *K_d_* is the apparent *K_d_*.

The equation 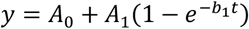 was used for fits to a single exponential function, where *A0* is the y-intercept, *A1* is the amplitude, and *b1* is the observed rate. For data fitting 3 exponentials, the following equation was used 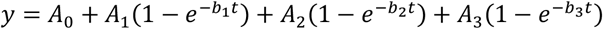 where *A0* is the y-intercept, *An* is the amplitude of the nth exponential and *bn* is the observed rate of the nth exponential.

#### 4.4.1. Data fitting by simulation with KinTek Explorer

Data fitting in KinTek Explorer was performed by fitting the kinetic data to one of the models given in the paper after providing the starting concentrations of reactants, initial estimates for rate constants, and an output observable. Confidence contour analysis was performed using the FitSpace (39) feature of KinTek Explorer. Parameter boundaries are reported in tables from kinetics experiments using a *χ*^2^ threshold in the FitSpace calculation which was recommended by the software as a reasonable limit based on the number of parameters and number of data points used in the fitting to give the 95% confidence interval. The threshold used is shown as the dashed grey lines in the confidence contours. Calculations of Flux in comparing fraction of reactions proceeding via alternative branched pathways were performed using a dynamic partial derivative analysis during data fitting based on numerical integration of the rate equations using KinTek Explorer software. The King-Altman method (44) was used to derive steady state kinetic parameters for each nucleotide incorporation pathway.

### 4.5. Molecular dynamics simulations

All atom MD simulations were performed on three systems comprised of T7 gene product 5, thioredoxin, a DNA primer/template of 27 or 28 and 45 bases, respectively, with an incoming 2xMg^2+^-nucleotide complex. The enzyme consists of 704 amino acids for gene product 5 and 109 amino acids for thioredoxin. The three systems studied are as follows:

1. <u>*T:dATP-Correct Nucleotide Complex:*</u> T7 DNA polymerase bound to a 27/45 nt primer/template DNA with a templating T, as well as an incoming dATP nucleotide and two Mg^2+^ ions.
2. <u>*T:dTTP-Mismatched Nucleotide Complex:*</u> T7 DNA polymerase bound to a 27/45 nt primer/template DNA with a templating T, as well as an incoming dTTP nucleotide and two Mg^2+^ ions. The structure from *T:dATP-Correct* was used as starting coordinates for MD simulation by replacing the dATP with dTTP.
3. <u>*A:dTTP-Mismatch Extension Nucleotide Complex:*</u> T7 DNA polymerase bound to a 28/45 nt primer/template DNA with a terminal T:T mismatch and a templating A, as well as an incoming dTTP and two Mg^2+^ ions. The structure from *T:dATP-Correct* was used as starting coordinates, after changing the DNA sequence to represent the DNA after T:T misincorporation with a terminal T:T mismatch and replacing dATP with dTTP as the incoming nucleotide.

#### 4.5.1. Initial structure preparation

The initial coordinates used to prepare all structures for MD simulations were based on the crystal structure of T7 DNA polymerase bound to DNA and ddATP (pdb:1skr (24)). All water molecules were removed and the missing residues were filled in with COOT (66). The DNA sequence was changed to match the sequence used for kinetics experiments (see Table 1). For that, the duplex portion of the DNA was extended, and the sequence was changed to match the DNA used in kinetics experiments, the single stranded template extension was assumed in a random conformation with no steric clashes. The terminal 2’,3’ dideoxy nucleotide of the primer strand was substituted with a base containing a 3’OH group and the incoming ddATP was replaced with dATP. The two Mg^2+^ ions bound to the incoming dATP were kept in their original positions.

#### 4.5.2. General MD simulation set up

All atom MD simulations were performed with GROMACS v5.0.7 (67). The models were immersed in a triclinic box with a minimum of 10 Å solvent edge from all directions. Explicit water and ions were added to mimic the experimental condition. Periodic boundary conditions were employed in all directions. To model water TIP3P (68) for other species amber14sb force field parameter (69) was employed. The number of cations were adjusted to mimic the concentration used in the kinetics experiments (12.5 mM MgCl_2_ and 50 mM NaCl respectively). Non-bonded interactions were truncated with a cut-off of 12 Å. Dispersion corrections were made to treat van der Waals interactions, the particle-mesh Ewald sum (PME) method was used for long range electrostatics.

The complexes were first energy minimized with 1,000 steps of steepest descent energy minimization to avoid bad contacts. This step was followed by volume equilibration using 2 ns long NPT simulations at 293 K and 1 bar using Berendsen thermostat (70). The LINCS algorithm (71) was used to constraints all covalent bonds. Next, we equilibrated the solvent (ions and water) by sampling conformations in NVT ensemble for 50 ns, keeping the temperature at 293 K with velocity scaling implemented in GROMACS. The equations of motions were integrated with a time step of 2 fs using Leap-Frog algorithm (72). During the equilibration, all non-hydrogen atoms of the complex were position restrained with a force constant of 1,000 kJ mol^−1^ nm^−2^ while cations and water molecules were allowed to move freely. The solvent equilibration step is followed by relaxing the single strand template region of the DNA. For that, we removed the position restraints for the single stranded region added while we kept other parts of the complex restrained. We sampled conformations for another 50 ns to relax this region. Last frame of the simulation was extracted and used to initiate an unrestrained production run. Equilibrium conformations were sampled for 500 ns in the NVT ensemble, keeping the simulation set up the same in the previous step. Coordinates were saved every 2 ps for data analysis.

#### 4.5.3. Preparation of T:dTTP mismatch and dTTP mismatch extension complexes

The T:dTTP mismatch complex, and the dTTP mismatch extension complex were prepared based on the T:dATP correct nucleotide complex listed above. The procedures for equilibration and production were as described above.

#### 4.5.4. Trajectory analysis

Trajectory analysis was performed using GROMACS suite of programs. A representative structure was selected for each complex when the structure figures are presented based on the average catalytic site distances obtained from each conformational pool. Molecular renderings were prepared with the PyMOL Molecular Graphics System, Version 2.0 Schrödinger, LLC (73). GROMACS analysis tools were also used for quantitative assessment of the trajectories given in Table 7.

## Data availability statement

All data are contained within the article.

## Acknowledgements

This work was supported by grants from NIGMS (5R01GM114223 to KAJ) and the Welch Foundation (F-1604 to KAJ). Computational research was carried out on the High Performance Computing resources at New York University Abu Dhabi. S.K. is supported by AD181 faculty research grant.

